# Allele-specific expression changes dynamically during T cell activation in HLA and other autoimmune loci

**DOI:** 10.1101/599449

**Authors:** Maria Gutierrez-Arcelus, Yuriy Baglaenko, Jatin Arora, Susan Hannes, Yang Luo, Tiffany Amariuta, Nikola Teslovich, Deepak A. Rao, Joerg Ermann, Helena Jonsson, Cristina Naverrete, Peter K. Gregersen, Tonu Esko, Michael B. Brenner, Soumya Raychaudhuri

## Abstract

Understanding how genetic regulatory variation affects gene expression in different T cell states is essential to deciphering autoimmunity. We conducted a high-resolution RNA-seq time course analysis of stimulated memory CD4^+^ T cells from 24 healthy individuals. We identified 186 genes with dynamic allele-specific expression, where the balance of alleles changes over time. These genes were four fold enriched in autoimmune loci. We found pervasive dynamic regulatory effects within six HLA genes, particularly for a major autoimmune risk gene, *HLA-DQB1*. Each *HLA-DQB1* allele had one of three distinct transcriptional regulatory programs. Using CRISPR/Cas9 genomic editing we demonstrated that a single nucleotide variant at the promoter is causal for T cell-specific control of *HLA-DQB1* expression. Our study in CD4^+^ T cells shows that genetic variation in *cis* regulatory elements may affect gene expression in a lymphocyte activation status-dependent manner contributing to the inter-individual complexity of immune responses.

## Main text

Genetic studies have identified an enrichment of autoimmune risk alleles in memory CD4^+^ T cell-specific regulatory elements(*1–3*). Memory CD4^+^ T cells are essential orchestrators of immune response. Hence, it is crucial to study how genetic variation affects their gene expression patterns to unravel the complex dynamics of regulation. Previous studies on activated T cells analyzed a limited number of cell states and genes(*4–9*), and an understanding of how gene expression levels are influenced by genetic regulatory variation in multiple physiological states is lacking. In this study, we investigated activation-dependent genetic regulatory effects in memory CD4^+^ T cells by studying dynamic allele-specific expression that changes with time in a high resolution RNA sequencing time series.

Studying allele-specific expression (ASE) of genes can enable the detection and characterization of context-specific *cis* regulatory effects (*10, 11*). In a pilot experiment, we stimulated memory CD4^+^ T cells from two genotyped individuals of European ancestry (fig. S1) with anti-CD3/CD28 beads. We ascertained gene expression at 0, 2, 4, 8, 12, 24, 48 and 72 hours after stimulation using deep mRNA sequencing (**Fig. 1A**). Using a logistic regression framework, we identified dynamic ASE (dynASE) events (**Methods**) at heterozygous SNPs. These dynASE sites are those where the imbalance of the two expressed alleles is time dependent. First, for each heterozygous site in an individual, we merged counts from all time points and identified 1,484 sites with evidence of significant ASE (intercept P < 2.8×10^−6^=0.05/17,743 tests, Bonferroni threshold). Next, for those sites we assessed time-dependent ASE effects by fitting a second order polynomial model. To account for over-dispersion of allelic counts, we incorporated sample-to-sample variability with a random intercepts effect. We observed 64 dynASE events in these two individuals (P < 3.7e-03, <5% FDR) in 60 SNPs, in 37 genes. In an independent experiment for the same two individuals, we observed that these dynASE sites had strong evidence of replication (~70% of events had P < 0.05), and a high correlation of effects for time (Spearman rho: 0.92 and 0.86) and time squared (Spearman rho: 0.51 and 0.68; fig S2–3).

**Fig. 1.**
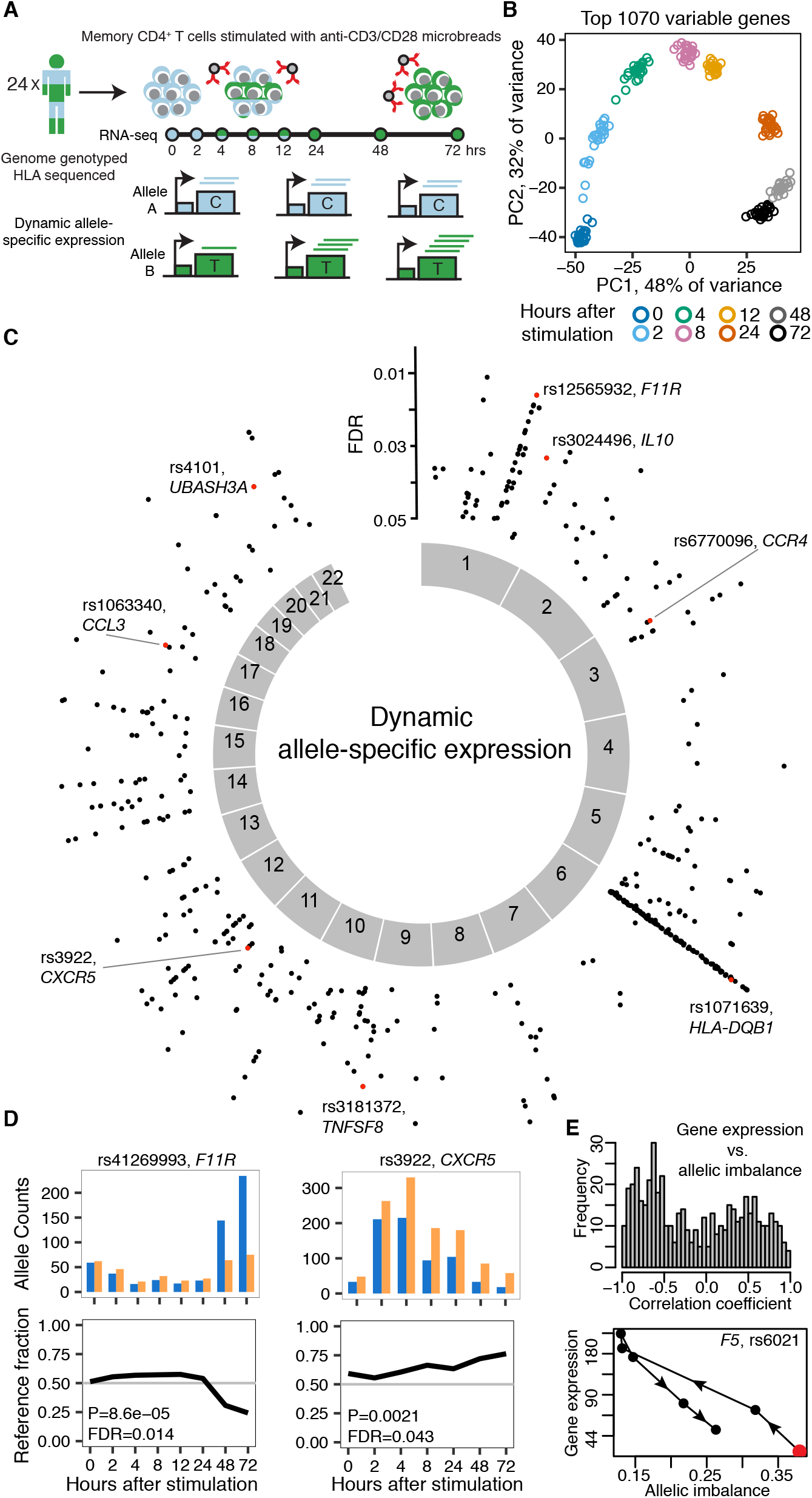
Dynamic allele-specific expression during T cell activation. (A) Study design. (B) Principal Component Analysis on top 1070 most variable genes. Shown are PC1 and PC2 scores for the 200 samples colored by time point. (C) Plot showing position across the genome of dynamic allele-specific expression (ASE) events, with y-axis indicating FDR. In red, highlighted examples. (D) Examples of dynamic ASE events in three genes. For each time point, we show allele counts for the SNP (top) and fraction of reads with the reference allele (bottom). (E) Spearman correlation coefficient between gene expression levels (in log2 scale) and SNP allelic imbalance (distance to 0.5 reference fraction) across time (top), and specific example (bottom). Red dot indicates start of trajectory (0 hour time point).

Next, we assayed an additional 22 individuals of European descent using the same experimental set-up to obtain data in a total of 24 healthy subjects (fig. S1). Principal component analysis on gene expression levels showed that the 200 samples separated by time (**Fig. 1B**, fig. S4), and gene clustering identified expected activation and repression clusters (fig. S5, **Methods**). To identify ASE, we queried a total of 225,924 events, representing 38,890 unique SNPs in 8,322 genes and some in transcribed intergenic regions (3%) (fig. S6). We observed a total of 15,268 ASE (P < 2.4e-07) events (2,147 genes). We then tested each of these events for dynASE and observed 561 significant events (P < 3.2e-03, <5% FDR), representing 356 SNPs in 186 genes and seven intergenic sites (**Fig. 1C**, Table S1). We found that 74% of our dynASE genes have been reported to have an eQTL in resting T cells (*4, 5, 8*), with an enrichment for T cell specific eQTLs (fig. S7). This indicates that we captured and expanded upon known *cis* regulatory genetic effects. **Fig. 1D** shows examples for SNPs in the genes *F11R* and *CXCR5*, where the reference and alternative alleles were dynamically regulated in time. These genes are critical for the migration of T cells across transendothelial membranes and within lymphoid tissues, respectively. Particularly, CXCR5 expression is important for T follicular helper cell localization in germinal centers. Interestingly, gene expression changes were coordinated with allelic imbalance changes over time (**Fig. 1E**). Hence, dynamic allelic imbalance suggests complex and continuous regulation during T cell activation affected by genetic variation. Consistent with this, dynASE genes were enriched for “immune response” function (P = 2.9e-09), even with respect to the genes with significant overall ASE (P = 2.5e-05, **Methods**).

Strikingly, we observed 182 dynASE events within the MHC locus (**Fig 1C**), with 15 events in *HLA-DQB1* (examples in fig. S8, Table S1), which harbors most of the genetic risk for type 1 diabetes (T1D) and celiac disease (*12, 13*). *HLA-DQB1* is part of the HLA class II genes, that are typically expressed in antigen presenting cells, and present antigen to CD4^+^ T cells to coordinate immune response. In human T cells, these genes are recognized as activation markers. Their expression is well characterized in T cells from the synovial fluid and tissue of rheumatoid arthritis patients, and in blood from patients with a variety of autoimmune disorders (*14–18*). Literature suggests that T cells themselves may also present viral and self-peptides to alter immune responses (*19–21*). However, further investigation is needed to confidently establish the function of HLA class II genes in T cells.

To better understand the relationship between HLA classical alleles (a combination of multiple coding variants) and regulatory variation, we performed high resolution typing of *HLA-DQB1, HLA-DRB1, HLA-DQA1, HLA-A, HLA-B*, and *HLA-C*, for all 24 individuals in our dataset (see **Methods**). To robustly identify dynASE in this highly polymorphic region, we built an HLA-personalized genome for each individual (**Fig. 2A**) by adding the cDNA allelic sequences for each HLA allele to the reference genome. We then quantified the number of uniquely mapped reads to each HLA allele per individual. Among the 48 *HLA-DQB1* ~780bp sequences, there were 14 *HLA-DQB1* 4-digit classical alleles. Our unique HLA-personalized genome strategy allowed us to quantify expression of individual *HLA-DQB1* alleles taking advantage of >20 SNPs in four exons (**Fig. 2B**, replication fig. S9). Our study underscores the importance of using a genotype-based personalized genome strategy to quanity *cis* regulatory effects within the HLA region.

**Fig. 2.**
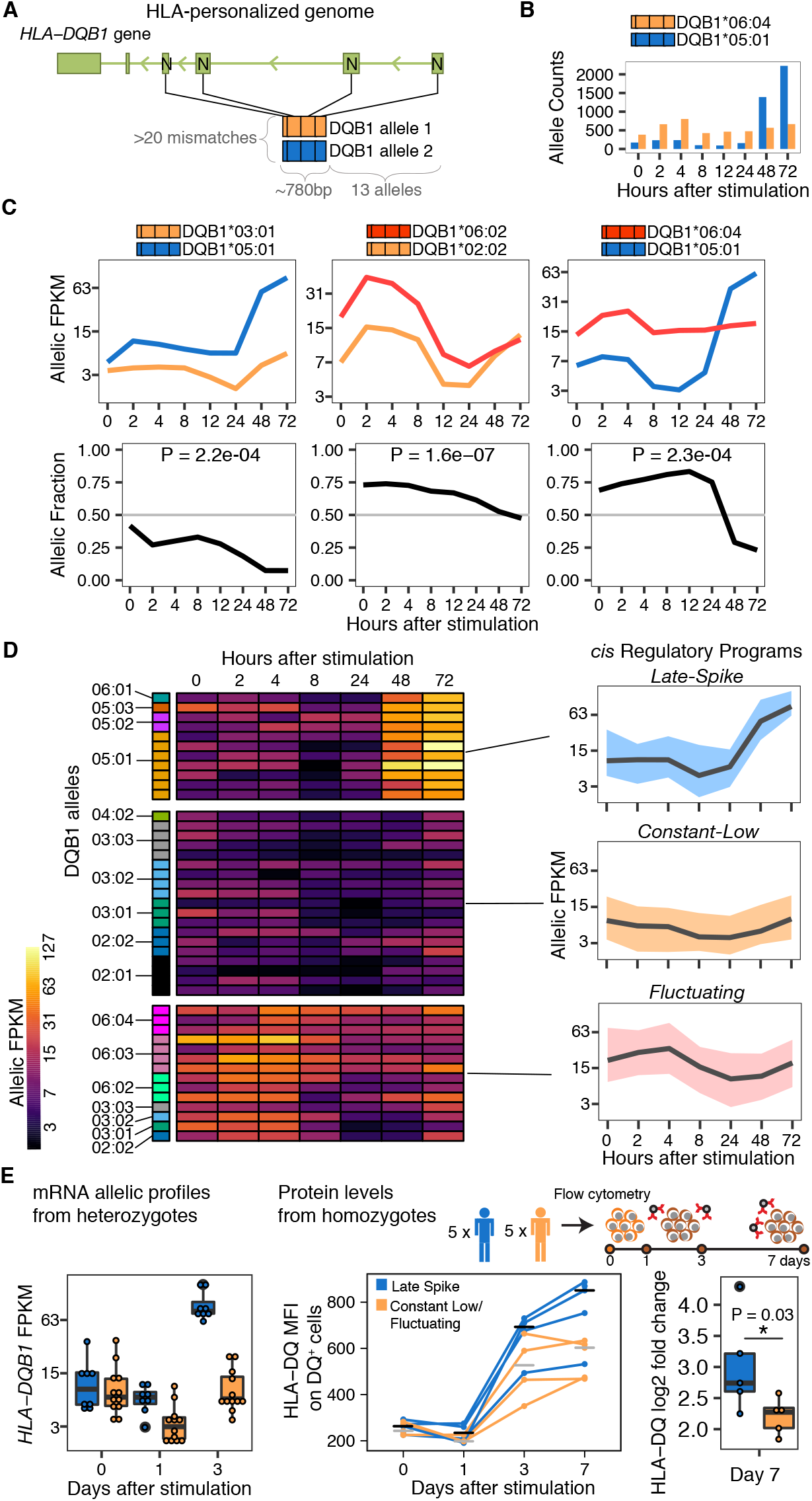
*HLA-DQB1* dynamic allele-specific expression at mRNA and protein levels. (A) Scheme of HLA-personalized pipeline and study design for allelic expression quantification in highly divergent *HLA-DQB1* alleles within each individual. (B) *HLA-DQB1* allele counts for an individual over time. (C) Normalized allelic expression for *HLA-DQB1* (top, in log2 scale), and allelic fraction (bottom) for three individuals. (D) Heatplot shows normalized allelic expression levels (in log2 scale) for each of the 48 HLA-DQB1 alleles in our cohort. Allelic profiles were clustered into three *cis* Regulatory Programs, for which the average expression profile is shown on the right with a black line, and total expression area occupied by all alleles in that cluster is shown with the colored ribbon. (E) Left panel shows normalized allelic mRNA expression levels (in log2 scale). Middle panel shows protein levels (median fluorescence intensity of HLA-DQ^+^ CD4^+^ memory T cells) for 5 homozygous individuals for alleles within the *Late-Spike* regulatory program (blue) and 5 homozygous individuals for alleles in *Constant-Low* or *Fluctuating* programs (yellow). Right panel shows log2 fold change in HLA-DQ MFI between day 0 and day 7. P-value from Wilcoxon one-tailed test.

Using allelic counts over the four exons of *HLA-DQB1*, we determined that most (15/24) individuals have significant dynASE for *HLA-DQB1* (P < 0.002 = 0.05/24 tests). For example, **Figure 2C** depicts profiles of allelic expression levels for three individuals, with their respective allelic fraction patterns over time.

Clustering of allelic expression profiles revealed that *HLA-DQB1* 4-digit classical alleles clustered together more than expected by chance (permutation P < 0.001, fig. S10), suggesting *cis* regulatory effects segregate with *HLA-DQB1* classical coding alleles. These allelic profiles could be grouped into three transcriptional groups (**Fig. 2D**, shown also with PCA on fig. S11). We named these three *HLA-DQB1 cis* regulatory programs based on their expression dynamics: *Late-Spike, Constant-Low*, and *Fluctuating* (**Fig. 2D**). To our knowledge, the identification of three distinct transcriptional profiles in *HLA-DQB1* expression over time is the first description of such complex and variable regulation in any gene.

To confirm that the drastic mRNA up-regulation in the *Late-Spike cis* regulatory program affected cell-surface protein expression levels, we isolated memory CD4^+^ T cells from an independent cohort. We specifically recruited five homozygous individuals for *HLA-DQB1* classical alleles with the *Late-Spike cis* regulatory program, and five homozygous individuals for *HLA-DQB1* classical alleles with *Constant-Low* or *Fluctuating cis* regulatory programs (fig. S12). As predicted, we observed that HLA-DQ cell surface expression on HLA-DQ^+^ cells was significantly higher in individuals with the *Late-Spike* haplotype (P = 0.03, Wilcoxon test, **Fig. 2E**, fig. S13).

We then sought to identify the genetic variant driving the *Late-Spike cis* regulatory program with genetic and epigenetic fine-mapping tools. We called SNP genotypes and 4-digit classical HLA alleles in 2,198 fully sequenced genomes (**Methods**). We identified SNPs in tight linkage disequilibrium (LD) with the *Late-Spike HLA-DQB1* classical alleles, that together represent what we henceforth call the *Late-Spike* haplotype (**Methods**). While most of the SNPs in highest LD with this haplotype (r ≥ 0.98) were within the *HLA-DQB1* gene (89%), only six were intergenic (fig. S14A). An eQTL analysis in the 24 individuals of our initial cohort (**Fig. 1**) confirmed that these 6 intergenic SNPs explained 76% of *HLA-DQB1* expression variance at 72 hours (P = 3.8e-08, fig. S14B). We assayed and identified open chromatin regions with ATAC-seq in memory CD4^+^ T cells after 72 hours of stimulation (**Methods**). In the *HLA-DQB1* locus, the highest peak was located at the promoter and overlapped with SNP rs71542466 (**Fig. 3C**, fig. S15). From published ChIP-seq data, this SNP overlapped promoter and enhancer associated chromatin marks in primary memory CD4^+^ T cells (**Fig. 3C**) (*22*), as well as binding peaks for HLA class II regulators: RFX5 (*23*) (in LCLs) and the co-activator CIITA (*24*) (B cells; fig S15).

**Fig. 3.**
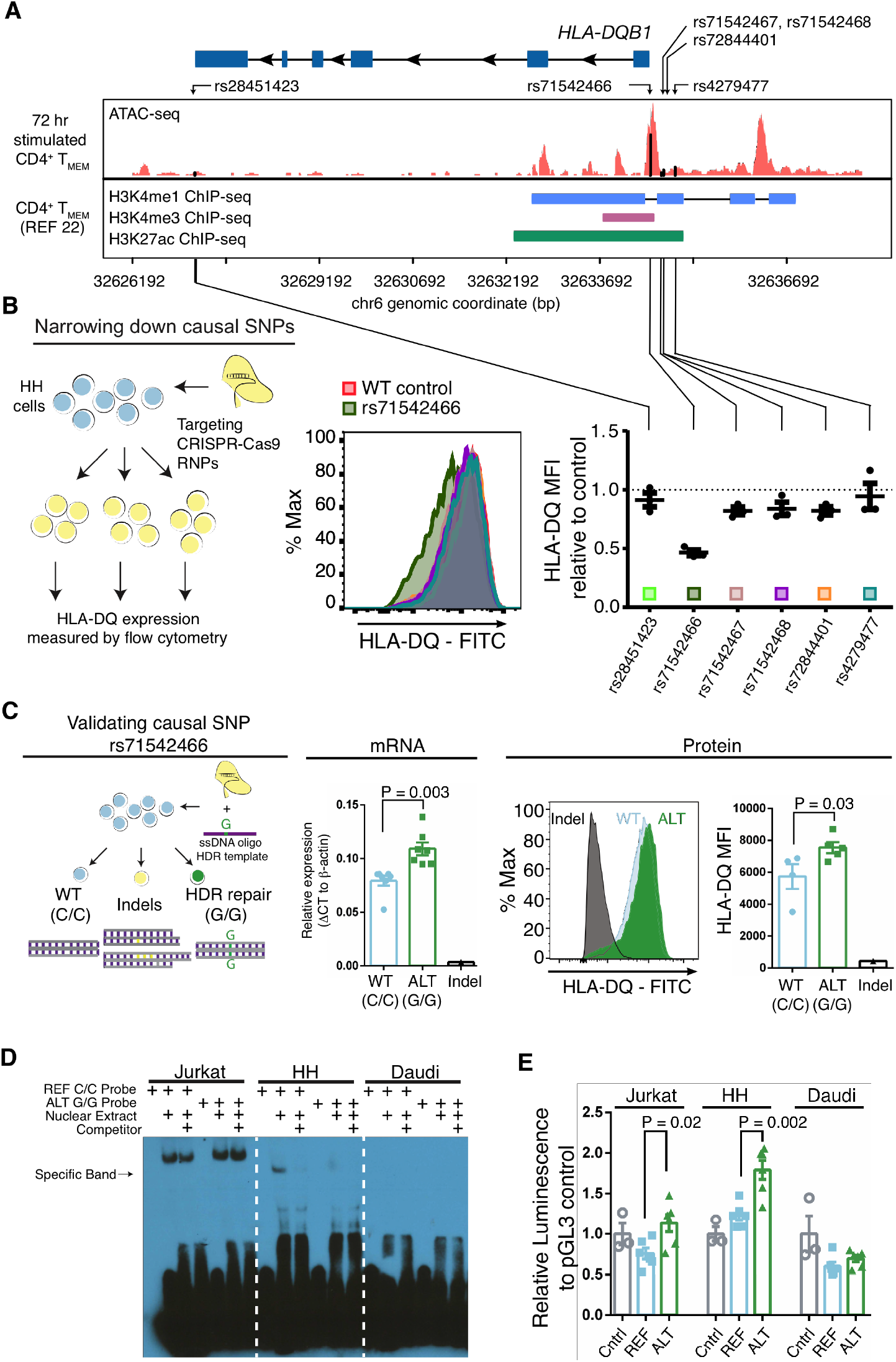
Validation of causal variant for *Late-Spike cis* regulatory program. (A) Location of 6 fine-mapped noncoding SNPs around *HLA-DQB1*. Tracks showing open chromatin regions (ATAC-seq) or regions marked by histone modifications (ChlP-seq). (B) CRISPR/Cas9 cuts at or near six fine-mapped SNPs in HH T cell lines. Left, experiment scheme. Middle, representative example of HLA-DQ expression levels. Right, HLA-DQ median fluorescence intensity relative to control, for each of the 6 SNPs, in triplicate. (C) Validation of causal SNP rs71542466 with CRISPR/Ca9 and ssDNA oligo HDR template. Left, experiment scheme. Middle, mRNA *HLA-DQB1* quantification with qPCR Taqman assay for 7 wild type (WT, CC genotype), and 7 SNP edited (ALT, GG genotype) expanded clones for rs71542466, as well as a cell line clone with an indel at the same target position. Right, HLA-DQ protein levels measured with flow cytometry. N = 5 WT and 4 ALT clones. (D) Electrophoretic Mobility Shift Assay using nuclear extract of three cell lines, with biotin labeled probes with reference (REF) or alternative (ALT) alleles for rs71542466. (E) Luciferase assay in three cell lines. Cntrl, control. REF, reference allele. ALT, alternative allele. All P-values from Mann-Whitney one-tailed test.

We hypothesized that amongst the 6 intergenic SNPs of the *Late-Spike* haplotype, rs71542466 located 39bp from the transcription start site of *HLA-DQB1* is causal. To test this, we employed CRISPR/Cas9 editing in the HLA class II expressing T cell line, HH (**Methods**). First, we designed guide RNAs to cut near the 6 SNPs (fig. S16A) in order to assess the regulatory potential of each region. We observed that only editing near rs71542466 caused a significant decrease in HLA-DQ expression (**Fig. 3D**, fig. S16B). Next, we applied targeted base-editing to rs71542466 in order to convert the reference C allele to the alternative G allele. We predicted that HH T cell line clones homozygous for the rs71542466 reference allele should have lower *HLA-DQB1* expression than base-edited clones with the alternative allele. We identified 7 clones homozygous for the alternative G allele, 7 clones homozygous for the reference C allele, and one clone with a 104bp deletion (fig. S17). As predicted, *HLA-DQB1* was higher in alternative clones as measured by real-time PCR (P = 0.003, **Fig. 3E**). We also noted that after extended culture, surviving alternative allele clones had higher expression of HLA-DQ protein as measured by flow cytometry (P = 0.03, **Fig. 3E**). These results confirmed that the rs71542466 promoter SNP changes *HLA-DQB1* expression in the expected direction in HH T cell lines, indicating that it at least partially accounts for the *Late-Spike cis* regulatory program.

After identifying the causal regulatory variant driving condition-dependent *HLA-DQB1* expression, we considered whether this effect is T cell-specific. We observed that our *Late-Spike* regulatory rs71542466 SNP (rSNP) was not in LD with reported eQTL SNPs for *HLA-DQB1* in B cell derived lymphoblastoid cell lines (LCLs), monocytes, and resting and infected macrophages (Table S2, r^2^≤0.27) (*5, 8, 25–27*). In fact, our rSNP had no association with expression of *HLA-DQB1* in resting macrophages (P = 0.81), in macrophages infected with Listeria (P = 0.31) or with Salmonella (P = 0.86) (*27*).

To confirm the function and specificity of the *Late-Spike* rSNP, we applied luciferase assays and Electrophoretic Mobility Shift Assays (EMSA) in three cell-lines; HLA-DQ negative Jurkat E6-1 T cells, and HLA-DQ positive HH T cells and Daudi B cells (fig. S18). EMSAs showed that only HH T cells had an allele specific band; we observed binding of nuclear extract proteins for the reference (C/C) but not the alternative (G/G) allele (**Fig. 3F**, fig. S19), suggesting that a repressor complex may bind to prevent upregulation of *HLA-DQB1*. Consistent with the base editing experiments, luciferase assays showed an increase in luminescence in the alternative (G/G) allele in HH T cells but not in Daudi B cells (**Fig. 3G**). Additionally, we stimulated primary B cells from the 5 homozygous individuals for the *Late-Spike* alleles and 5 homozygotes for the *Fluctuating* or *Constant-Low* alleles and did not observe fold change differences in HLA-DQ cell surface expression at 24 hours (P = 0.3, Wilcoxon test, fig. S20).

Intriguingly, within the HLA, dynamic gene regulation was not unique to *HLA-DQB1*. We observed dynASE in 5 other HLA genes. Within HLA class II genes, *HLA-DRB1* had significant dynASE in 9 individuals (P < 2.1e-03, Bonferroni threshold), and *HLA-DQA1* in 6 (P < 2.2e-03, Bonferroni threshold). Within HLA class I genes, *HLA-B* and *HLA-C* had significant dynASE in 12 individuals (P < 2.4e-03 and P <2.5e-03, Bonferroni threshold, respectively), and *HLA-A* in 6 (P < 2.2e-03, Bonferroni threshold). However, for these 5 genes, the magnitude of change in allelic fraction across time was more modest than for *HLA-DQB1* (fig. S21). We observed that in a subset of *Late-Spike HLA-DQB1* haplotypes, the *HLA-DRB1* allele that is in phase on the same chromosome also followed a *Late-Spike* pattern of expression (fig. S22A). This suggests potential promoter interactions between *HLA-DQB1* and *HLA-DRB1*, consistent with promoter capture HiC data (fig. S22B) (*28, 29*).

We found that 31 of our dynASE genes outside of the MHC were within autoimmune disease loci, including *UBASH3A* and *IL10* (**Fig. 4D**). We evaluated whether dynASE genes were significantly enriched in autoimmune disease loci using a stringent strategy where we compared the number of dynASE genes observed within risk loci and those found in 1000 null sets of loci across the genome (**Methods**). We found that dynASE genes are significantly enriched for risk loci for ankylosing spondylitis (OR = 5.7, P = 0.008), celiac disease (OR = 5.4, P = 0.004), vitiligo (OR = 5, P = 0.004), type 1 diabetes (OR = 4.5, P = 0.002), inflammatory bowel disease (OR = 3.7, P = 0.001), rheumatoid arthritis (OR = 3.6, P = 0.005), and multiple sclerosis (OR = 3.1, P = 0.003), but not for non-immune mediated diseases (**Fig. 4E**). We compared these enrichments to non-dynASE events prior to stimulation (those with ASE at 0 hours only) and to published eQTLs for resting CD4^+^ naïve T cells (*8*). We observed that our dynASE genes, spanning up to 8 different cellular states, had the highest enrichment for autoimmune disease genes (**Fig. 4F**). We found similar results when we assessed enrichment of polygenic heritability in the regions around these genes versus the rest of the genome using stratified LD score regression (fig. S23).

**Fig. 4.**
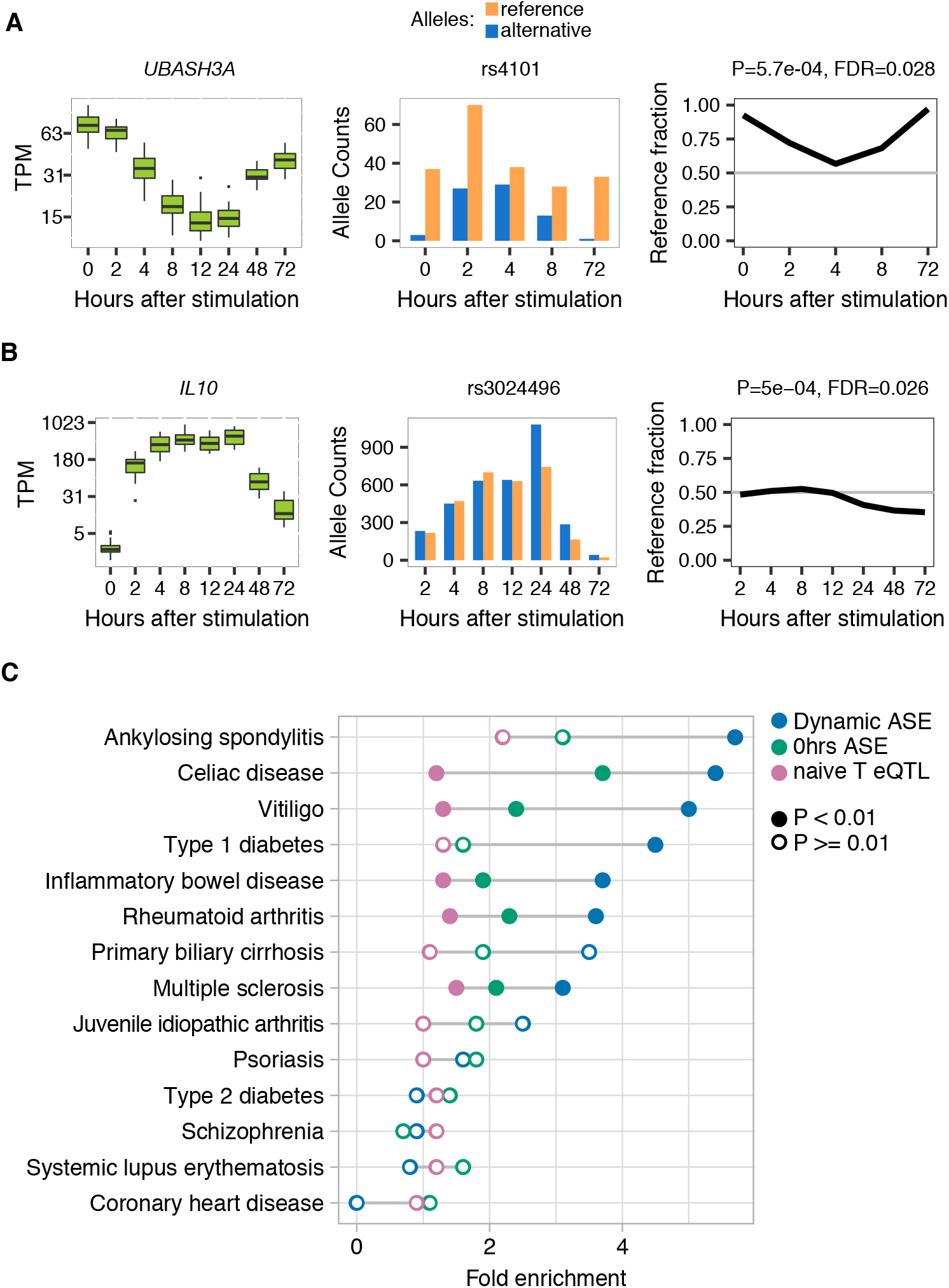
Non-MHC dynamic allele-specific expression genes are enriched in autoimmune disease loci. (A-B) DynASE examples for two autoimmune disease genes. (C) Fold enrichment of dynASE genes (blue) in risk loci for autoimmune diseases and 3 non-immune mediated diseases, using 1000 null sets of loci matched by number of loci per disease and number of genes per locus. Same for genes with significant ASE at 0 hours (green), and naive T cell eQTL genes (pink). Filled circles mark permutation P < 0.01, empty circles mark permutation P > 0.01.

In this study, we showed that allelic imbalance in expression is highly context-dependent and very sensitive to the time after stimulation of memory CD4^+^ T cells. In most cases, there was a dominant allele that gradually increased or decreased its expression dominance with time (e.g. *UBASH3A, CXCR5*). We suspect that these cases are a consequence of a single regulatory complex interacting with a single genetic variant altering gene expression; where the status of the regulatory complex may vary dependending on the environmental context. However, in many instances, we observed that the allele that was dominant switched over time (e.g. *HLA-DQB1, F11R*), which raises the possibility of multiple regulatory variants or complexes. For example, we identifed distinct driving variants for *HLA-DQB1* at 0 and 72 hours after stimulation (LD r^2^=0.01, Table S2). Overall, this widespread dynamic allelic imbalance across the genome illustrates the continuously changing regulatory landscape of genes during T cell activation.

We found that dynASE genes were highly enriched in autoimmune disease loci. This indicates that for many autoimmune disease genes, the risk allele may affect the expression of its target gene under very specific conditions. Indeed, this has already been shown for an autoimmune risk variant in an enhancer regulating *IL2RA* that affects its expression in a time-dependent manner, and the delay of expression by the risk allele leads to a preferential polarization of T cells into an inflammatory subtype (Th17) instead of the regulatory population (Treg) (*30*). These results may explain why investigators have found limited shared genetic effects between autoimmune susceptibility variants and eQTL variants at resting state (*3, 7, 31*), despite the presence of autoimmune susceptibility variants in regulatory elements (*1–3, 32*).

Intriguingly, we identified the most dramatic T cell and condition dependent *cis* regulatory variation within a major autoimmune disease gene: *HLA-DQB1*. This raises the question of whether, and to what extent, genetic regulatory variation controlling HLA gene expression could affect disease susceptibility or disease penetrance, as has been highlighted for other loci and traits (*33*). For most autoimmune diseases, the MHC region is the major contributing locus to disease risk. We and others have shown that specific amino acid changes in the peptide binding grove of *HLA-DQB1* and *HLA-DRB1* explain the majority, albeit not all, of the risk assigned to the MHC region in T1D and rheumatoid arthritis (*12, 34*). In this study, three of the four *Late-Spike HLA-DQB1* classical alleles are protective for T1D (OR 0.045-0.732), while the other two regulatory programs represent a mixture of risk and protective alleles (*12*). Detailed analyses on large sample sizes will be needed to disentangle the regulatory effects from the strong amino acid effects. While amino acid changes causing differential antigen display may be the primary autoimmune mechanism at the HLA locus, our data underscores the possibility that expression levels of HLA class II may also play a crucial and unappreciated role (*35, 36*). Over the past several decades, there has been literature suggesting variation in expression among different HLA alleles (*26, 37–39*) – but to date the idea that this regulation changes with cell-state has not been established. It is well known that positive selection has resulted in dramatic coding variation within most HLA genes. Our results suggest that the same positive selection may also have favored regulatory variation in *HLA-DQB1* and other HLA genes.

Broadly, class II expression has been well-characterized as a marker for both late activation in CD4^+^ T cells and suppressive capacity in T regulatory cells (*40–42*). However, the exact mechanisms and functional implications remain to be defined. Our work shows that not only do CD4^+^ T cells express high levels of HLA Class II, but that its expression is regulated in a cell-type specific manner and varies between individuals. This suggests that during immune responses, expression of HLA II on CD4^+^ T cells is dynamically controlled and may be important to modulating function.

Overall, by using both computational and experimental tools, we identified and validated highly context-specific genetic regulatory effects in stimulated memory CD4^+^ T cells and confirmed a causal variant in a key autoimmune gene. This work provides a framework for future studies to identify and validate relevant disease genes and variants.

## Methods

### Study design

The goal of this study was to characterize cell state-dependent regulatory effects in memory CD4^+^ T cells to obtain new insights into autoimmune disease mechanisms. In initial experiments, we observed the most dramatic dynamic allele-specific expression effects in *HLA-DQB1*. To investigate this phenomenon in an optimal way, we recruited individuals heterozygous for common *HLA-DQB1* classical alleles (characterized by a combination of coding single nucleotide alleles), with highly divergent alleles (at least a 20 mismatch differences between the two alleles). We used the de-identified genome-wide genotypes available from the individuals at the Genotype and Phenotype (GaP) Registry at The Feinstein Institute for Medical Research, to impute HLA classical alleles with SNP2HLA (*43*) and select individuals. The GaP Registry provided de-identified cryopreserved PBMCs from 24 donors with no autoimmune disease, 20-50 years old, and of European ancestry. Donors provided fresh, de-identified human peripheral blood mononuclear cells (PBMCs); blood was collected from subjects under an IRB-approved protocol (IRB# 09-081) and processed to isolate PBMCs. The GaP is a sub-protocol of the Tissue Donation Program (TDP) at Northwell Health and a national resource for genotype-phenotype studies (*44*). HLA classical alleles were subsequently experimentally confirmed with HLA typing (see below).

Similarly, for the protein level validation experiments, we recruited through the GaP Registry individuals homozygous for *HLA-DQB1* classical alleles pertaining to the *Late-Spike* regulatory program (N = 5) or other programs (N = 5). These individuals were also between 20 to 50 years old, with no reported autoimmune disease, and of European ancestry.

### Memory CD4^+^ T cell stimulation time course

PBMCs were thawed and resuspended in pre-warmed complete RPMI (cRPMI) (RPMI 1640, supplemented with 10% heat inactivated FBS, and 1% non-essential amino acids, sodium pyruvate, HEPES, L-Glutamine, Penicillin & Streptomycin, and 0.1% β-mercaptoethanol). Memory CD4^+^ T cells were isolated by magnetic selection (Miltenyi, Memory CD4^+^ T cell Isolation Kit human). Cells were counted, washed in cRPMI, and resuspended at a concentration of two million cells per ml. One million cells per well were plated in sterile 48 well plates (Corning). The cells were rested at 37°C overnight. Twelve hours after the beginning of the rest marked the first time point, T = 0 hours. At this time, cells were spun down and resuspended in 350 μL of RLT lysis buffer (Qiagen) containing β-mercaptoethanol and stored in a sterile Eppendorf tube at −80°C. To the remaining wells, 500 μL cRPMI with human T-Activator CD3/CD28 Dynabeads (Gibco) were added to each well at a ratio of 2 cells: 1 bead. Cells were collected at 2, 4, 8, 12, 24, 48, and 72 hours after stimulation. Once all cell pellets were collected, resuspended in RLT and frozen, the mRNA was isolated using a RNeasy mini kit (Qiagen). RNA concentration was measured using Implen’s cuvette based UV/Vis spectrophotometer. The purified RNA samples were stored at −80°C until submitted for sequencing at the Broad Institute in Cambridge, MA.

### Library construction and RNA sequencing

Total RNA was quantified using the Quant-iT™ RiboGreen^®^ RNA assay kit and normalized to 5 ng/μL. Following plating, 2 μL of ERCC controls (using a 1:1000 dilution) were spiked into each sample. An aliquot of 200 ng for each sample was transferred into library preparation which uses an automated variant of the Illumina TruSeq™ stranded mRNA sample preparation kit. Briefly, oligo dT beads were used to select mRNA from the total RNA sample followed by heat fragmentation and cDNA synthesis. The resultant cDNA was then dual-indexed by ‘A’ base addition, adapter ligation using P7 adapters, and PCR enrichment using P5 adapters. After enrichment, libraries were quantified using Quant-iT PicoGreen (1:200 dilution). After normalizing samples to 5 ng/μL, the set was pooled and quantified using the KAPA library quantification kit for Illumina sequencing platforms. The entire process was performed in 96-well format and all pipetting performed by either Agilent Bravo or Hamilton Starlet.

Pooled libraries were normalized to 2 nM and denatured using 0.1 M NaOH prior to sequencing. Flowcell cluster amplification and sequencing were performed according to the manufacturer’s protocols using either the HiSeq 2000 or HiSeq 2500. Each run used 101 bp paired-end reads with eight-base index barcodes. Data was de-multiplexed and aggregated using the Broad Picard pipeline. Libraries were sequenced at a mean depth of 41 million fragments (read pairs), median 37 million, and minimum 24 million fragments.

### Gene expression analyses

We mapped reads to the hg19 reference genome with subread v1.5.1(*45*) (with parameters: -u -Q -D 100000 -t 0 -T 4) and quantified expression levels using featureCounts (with parameters: -T 4 -Q 20 -C -s 2 -p -P -D 100000) and GENCODE (*46*) v19 annotation. We removed two outliers, one had >50% of PCR duplicates and <60% of reads assigned to genes, the other one had less than 19,000 genes detected and less than 98% of common genes detected (common genes are those detected in at least 95% of samples; fig. S4). We considered expressed genes with log2(tpm+1) > 2 in at least 20 samples. For PCA we took 1,070 genes with standard deviation > 1 and mean expression > 3 log2(tpm+1), we scaled gene to mean zero and variance one and performed PCA with the R (*47*) function prcomp. For gene clustering, we used k-means on the scaled mean expression per gene. Six clusters captured 70% of between group over total sum of squares; increasing k, the predefined number of clusters, had minor incremental improvements. Enrichment of MsigDB hallmarks v6.2 (*48, 49*) was performed with the enricher function of the R package clusterProfiler (*50*).

### Variant genotyping, imputation and filtering

Individuals were genotyped genome-wide using the Genome Screen Array (GSA) assaying 647K SNPs. For pre-imputation QC, we used plink v1.90b3w (*51*) to filter out variants with missing call frequencies greater than 0.05, Hardy Weinberg Equilibrium (HWE) threshold P < 1e-05, MAF 0.03, keeping a total of 339,333 variants. We imputed variants into the 1000 Genomes reference panel (*52*) using SHAPEIT v2.r837 (*53*) and IMPUTE2 v2.3.2 (*54*). We filtered out variants with info score < 0.9, multiallelic, HWE threshold P < 1e-05, non-polymorphic within our 24 individuals, and with MAF <1% in Europeans of the 1000 Genomes reference panel, and indels. This yielded a total of 5,144,453 SNPs. When selecting heterozygous SNPs per individual, we further required a genotype probability > 0.9; a total of ~1.5M heterozygous SNPs per individual remained.

### Dynamic allele-specific expression analysis

We used subread v1.5.1 to align reads to the hg19 reference genome and filtered out reads with mapping quality < 10. We used WASP (*55*) to filter out reads that had mapping bias at heterozygous sites and to remove duplicates. For quantifying allele counts at heterozygous sites, we used GATK (*56*) v3.8 ASEReadCounter (with parameters --minMappingQuality 10, -- minBaseQuality 10, -U ALLOW_N_CIGAR_READS), and followed recommended best practices (*57*). For initial QC (fig. S6), for each sample we took all heterozygous sites with at least 10 reads (6,496-44,864 sites per sample). The mean coverage across sites per sample ranged from 50-127. All samples had >95% of both alleles observed at included heterozygous sites. The mean reference fraction was close to 0.5 for all samples (mean 0.5098, range from 0.5045 to 0.5159). For a given heterozygous site, the reference fraction refers to the number of reads with the reference allele divided by the total number of reads overlapping the site. Allelic imbalance is the distance to 0.5 reference fraction (i.e. absolute value of: the reference fraction minus 0.5).

To identify sites with dynamic ASE, we used a nested approach using a logistic regression framework on a per individual, per heterozygous site basis, with the *lme4* (*58*) R package. Each read is encoded according to the following: 1 if it contains a reference allele or 0 if it contains the alternative allele. For each time course per individual, we included sites with at least 20 reads in at least 4 time points and required that both reference and alternative alleles are seen in all included samples. First, we identified sites with evidence of ASE by merging data from all time points and testing if the intercept is significantly different from zero (assuming a standard normal distribution and using a z-test) and used a Bonferroni threshold to determine significance (0.05 divided by the number of tests). Then, we tested which of these sites had ASE that changes with time by fitting a second order polynomial model, coding time point 1 through 8 (or maximum number of time points) and scaling to mean zero variance one. We controlled for overdispersion of allelic counts due to technical or biological sample to sample variability (*59*) by incorporating a random intercept effect, coding sample ID as a factor. We tested for the effects of time by performing a likelihood ratio test between the two nested models using R anova function. The null model *H*_0_ and alternative model *H*_1_ for a given SNP *i* in a given individual *j* are detailed below:

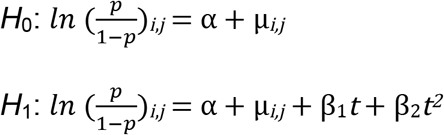

Where *p* is the probability of observing the reference allele, α is the intercept, μ is the random intercept effect across samples, *t* is time, and β is the effect of time on the log-odds of observing the reference allele. We calculated the FDR per test using the *qvalue* R package (*60*), and called significant all events with q<0.05 (<5% FDR), unless otherwise stated.

Functional enrichment for dynASE genes was performed using the C5 genesets from MsigDB, which are composed of GO biological processes (c5.bp.v6.2). Enrichment with respect to all tested genes or all genes with significant ASE (over all merged time points) was tested with the enricher function, of the clusterProfiler R package, which is based on the hypergeometric distribution.

### HLA typing

Purified DNA from the 24 individuals was sent to the NHS Blood and Transplant, UK, where HLA typing was performed. Next generation sequencing was done for *HLA-DQB1, HLA-A, HLA-B, HLA-C*, and for *HLA-DRB1*. PCR-SSOP was done for HLA-DQA1 in all individuals, and for HLA-DRB1 in 6 individuals for which limited DNA was available. These typing methods yielded classical allele calls for the six genes at 4 to 8-digit resolution.

### Allele-specific expression in HLA genes

To prepare an HLA personalized genome for each individual we first took the HLA 6-digit classical allele calls for each of the 6 HLA genes (12 alleles total) and downloaded from the IPD-IMGT/HLA database (*61*) the corresponding cDNA sequence. Next, we added the 12 cDNA sequences to the hg19 reference genome, each encoded as a separate chromosome. We masked with Ns the exonic regions corresponding to the cDNA sequences. We indexed the genome for subread usage.

For each individual, we aligned per sample the reads to the HLA-personalized genome with subread (with parameters: -u -Q -D 100000 -t 0 -T 4). We removed PCR duplicates with Picard Tools v1.119. We counted the number of uniquely mapped reads to each HLA allele with featureCounts (with parameters: -T 4 -Q 40 -p) using a personalized gtf annotation file per individual.

To identify dynASE for each HLA gene, we used the same statistical approach mentioned above. Instead of using allele counts for a single SNP, we used counts for the whole cDNA per HLA allele (usually encompassing 3-4 exons, 552-1119 bp). To compare HLA allelic expression levels between samples, we normalized the HLA allele counts by library size and cDNA size (FPKM). For the two individuals for which we had full time course replicates, we used the mean of the FPKM values.

For testing whether allelic profiles of *HLA-DQB1* 4-digit classical allele groups are more similar to each other than expected by chance, we calculated the sum of squares within 4-digit allele groups, and total sum of squares (observed values). Then, we permuted the 4-digit allele groups 10,000 times, and repeated the sum of squares calculations (fig. S10). For this analysis, we excluded four allelic profiles with 4-digit alleles that were present only once in our 48 allelic profiles (DQB1*04:02, DQB1*06:01, DQB1*06:09 and DQB1*05:03).

For *HLA-DQB1* allelic profile grouping with k-means clustering and PCA, we removed the 12 hour time point due to its high number of missing values (caused by an insufficient number of cells obtained for some individuals), and further excluded another 2 individuals with missing values, resulting in a total of 44 allelic profiles (2×22 individuals). For both of these analyses we used log2(FPKM+1) values. Three clusters captured 64% of the variation in our data, defined by the ratio of between group sum of squares and total sum of squares. Two clusters captured a significantly lower amount of between group variability (37%), and four clusters (70%) had a modest increase from three. PCA was performed on allelic expression profiles with prcomp with center = TRUE, and independently showed that the three clusters identified with k-means separate well (fig. S11).

### HLA-DQ protein level validations on memory CD4^+^ T cells and B cells

Ten additional individuals were recruited through the GaP Registry (see Study Design). Five individuals were homozygous for *Late-Spike* alleles: one for DQB1*05:03 and four for DQB1*05:01. Five individuals were homozygous for alleles in the other two *cis* regulatory programs (*Constant-Low*, and *Fluctuating*): one for DQB1*03:02, one for DQB1*02:01, one for DQB1*02:02 and two for DQB1*03:01. PBMCs were thawed and put immediately in warm cRPMI. For analysis of memory CD4^+^ T cells, cells were washed twice and isolated as before using a magnetic selection kit (Miltenyi, Memory CD4^+^ T cell Isolation Kit human). Isolated cells were then rested overnight at a concentration of 1 million/mL in 48 well plates and stimulated the next day with anti-CD3/CD28 dynabeads at a ratio of 2 cells: 1 bead. T cells were assayed for expression of HLA-DQ by flow cytometry on day 0 (unstimulated) and days 1, 3, and 7. For all samples, cells were isolated, washed twice in PBS, and stained with Zombie Violet Live/Dead (Biolegend) for 30 minutes on ice. After staining, cells were washed and Fc receptors blocked with FcX True Stain (Biolegend) for 15 minutes on ice followed by staining with directly conjugated antibodies: PE anti-HLA-DR (Biolegend, Clone: L243), APC anti-HLA-DP(Leinco Technologies, Clone: B7/21), FITC anti-HLA-DQ(Biolegend, Clones: HLADQ1 or Tu169), BV711 anti-CD25(Biolegend, Clone: M-A251), and PE-Cy7 anti-CD4(Biolegend, Clone: OKT4). Cell were stained for 30 minutes on ice, washed twice, and samples analyzed on a BD LSR Fortessa. For B cells, 0.5 million total PBMCs in cRPMI were stimulated with a cocktail of CpG (2006, 0.35 μM), CD40L (1 μg/mL), anti-Ig (10 μg/mL), and rhIL-21 (20 ng/mL) in 48 well plates for 0 and 1 day. As before, all samples were isolated, stained with Live/Dead, and Fc receptors blocked. Cells were then stained for 30 minutes on ice with PE anti-HLA-DR (Biolegend, Clone: L243), APC anti-HLA-DP (Leinco Technologies, Clone: B7/21), FITC anti-HLA-DQ (Biolegend, Clones: HLADQ1 or Tu169), PeCy7 anti-IgD (Biolegend, Clone: IA6-2), BV510 anti-CD27 (Biolegend, Clone: O323), BV605 anti-CD3(Biolegend, Clone: OKT3), BV785 anti-CD38 (Biolegend, Clone: HB-7), BV650 anti-CD19(Biolegend, Clone: HIB19), PerCP-Cy5.5 anti-CD45R0 (Biolegend, Clone: UCHL1), BUV496 anti-CD4 (BD Biosciences, Clone: SK3). As before, data was collected on a BD LSR Fortessa. All data was processed using Flowjo, gated as shown in the supplement, and channel values exported for analysis.

### Regulatory variant fine-mapping

To look for genetic variants in LD with the *Late-Spike* haplotype, we called Single Nucleotide Variants (SNVs), small INsertions and DELetions (INDELs) and classical HLA variants using whole genome sequences of 2,244 healthy volunteers recruited from the Estonian Genome Project (*62, 63*) sequenced at 25x. We performed high-resolution (G-group) HLA calling of three class-I HLA genes (HLA-A, -B and -C) and three class-II HLA genes (HLA-DRB1, -DQA1 and -DQB1) using the HLA*PRG algorithm (*64*). We called SNVs and INDELs using GATK version 3.6 according to the best practices for variant discovery (*65*). In total we called 246,505 variants in the extended MHC region (29-34 Mb on chromosome 6, NCBI Build 37).

To check if any SNVs are in high LD with the *Late-Spike* haplotype, we first used the Estonian reference panel and restricted our analyses to individuals who carried the alleles present in our 24 individuals (N = 2,198). Namely, there were 58 individuals with two *HLA-DQB1* alleles (at 4-digit resolution) pertaining to the *Late-Spike* haplotype (i.e. DQB1*05:01, 05:02, 05:03 or 06:01), 616 with one *Late-Spike* haplotype allele, and 1,524 individuals with zero *Late-Spike* haplotype alleles. To filter out possible false-positive variants, we next restricted the analyses to SNPs with minor allele frequency (MAF) ≥ 0.05 and within 1 Mb region of the *HLA-DQB1* gene (N = 27,210). Next, we computed the Pearson correlation between *Late-Spike* haplotype dosage (0, 1 or 2) and individual SNP genotypes. Refseq gene annotations were used to determine start and end of *HLA-DQB1, HLA-DRB1* and *HLA-DQA1*.

For the eQTL analyses within our cohort we used log2(FPKM+1) gene expression values for *HLA-DQB1* (the sum of FPKM of both alleles from each individual) at a given time point and looked for association of expression with SNPs (with MAF 5%) within 1Mb of the TSS using Pearson correlation.

### ATAC-seq experiments and data processing

Memory CD4^+^ T cells from one new PBMC donor from the GaP Registry were purified and cultured as described above for 72 hours with anti-CD3/CD28 stimulation beads. Post stimulation, the cells were harvested and washed with PBS. Cells were then resuspended in 500μL of a freshly prepared 1% formaldehyde solution (Thermo Scientific) and fixed for 10 minutes. The fixation reaction was subsequently quenched with the addition of 2.5 M glycine for 5 minutes. The sample was spun to remove the fixative and washed in PBS. The solution was carefully decanted and the resulting pellet was frozen. For open chromatin library preparation, two samples, one of 50,000 cells and another of 10,000 cells, were resuspended in 1 mL of cold ATAC-seq resuspension buffer (RSB; 10 mM Tris-HCl pH 7.4, 10 mM NaCl, and 3 mM MgCl_2_ in water). Cells were centrifuged at max speed for 5 min in a pre-chilled (4°C) fixed-angle centrifuge. After centrifugation the supernatant was carefully aspirated. Cell pellets were then resuspended in 50 μL of ATAC-seq RSB containing 0.1% NP40, 0.1% Tween-20, and 0.01% digitonin by pipetting up and down three times. This cell lysis reaction was incubated on ice for 3 min. After lysis, 1 mL of ATAC-seq RSB containing 0.1% Tween-20 (without NP40 or digitonin) was added, and the tubes were inverted to mix. Nuclei were then centrifuged for 5 min at max speed in a pre-chilled (4°C) fixed-angle centrifuge. The supernatant was removed and nuclei were resuspended in 50 μL of transposition mix (*66*) 2.5 μL transposase (100 nM final), 16.5 μL PBS, 0.5 μL 1% digitonin, 0.5 μL 10% Tween-20, and 5 μL water) by pipetting up and down six times. Transposition reactions were incubated at 37°C for 30 min in a thermomixer with shaking at 1,000 rpm. Reactions were cleaned up with Qiagen columns. Libraries were amplified as described previously (*67*).

Libraries were paired-end sequenced with an Illumina NextSeq 500 with read length of 150 bp. Reads were mapped to the hg19 reference genome with subread. Uniquely mapped reads with mapping quality > 10 were selected, and PCR duplicates were removed with Picard tools. Peaks on chromosome 6 were called with MACS2 (with parameters --nomodel -q 0.1).

### Cell lines

HH cutaneous T cell lines (ATCC: CRL-2105), Jurkat E6-1 (ATCC: TIB-152), and Daudi (ATCC: CCL-213, provided by Dr. Michael Brenner), were all cultured in complete RPMI as previously described.

### Bulk CRISPR/Cas9 editing

To investigate regulatory regions around HLA-DQ, the nearest sgRNA to the SNP of interest was selected using Deskgen online tools (www.deskgen.com). Distances of designed sgRNA to the nearest SNP are shown in fig. S16. To confirm the sequence of the region, genomic DNA around *HLA-DQB1* was PCR amplified and Sanger sequenced using the primers: 5’-TCGGGTCTCTGAATCCCACT & 5’CAGGACCCAGGAAATGCTTCT, and 5’GGAGCTCTGCCATTTGTCCT & 5’TGACTCTGCTTCCTGCACTG. CRISPR/Cas9 RNP complexes were assembled as previously described(*68*). Briefly, 40 μM Cas9 protein (QB3 Mircolabs) was mixed with equal volumes of 40 μM modified sgRNA (Synthego) and incubated at 37°C for 15 minutes to form ribonuclear protein (RNP) complexes. HH cells were nucleofected with 2μL of RNPs in an Amaxa 4D nucleofector (SE protocol: CL-120). Cells were immediately transferred to 24 well plates with pre-warmed media and cultured. After 7-10 days, HLA-DQ expression was assessed by flow cytometry. Editing was confirmed by PCR amplifying genomic DNA around *HLA-DQB1* and sequences analyzed by Tracking of Indels by Decompostiion (TIDE) analysis (tide.deskgen.com).

### sgRNA target sequences with PAM bolded

rs28451423 - TGTGAAATCAACTTGACTCT**AGG**,

rs71542466 - GCTGATTGGTTCTTTTCCGA**GGG**,

rs72844401 - AATGCCTCGGGGATTTTGAG**AGG**,

rs4279477 - AGAACTTTGCTCTTCTCCCC**AGG**,

rs71542467 - GAGCTGAAGAACGAATGCCT**CGG**,

rs71542468 - GCTGAAGAACGAATGCCTCG**GGG**

### CRISPR/Cas9 base-editing of rs71542466 in HH cells

For generation of base-edited cell lines, HH cells were nucleofected with RNPs using sgRNA targeting near rs71542466 and asymmetrical ssDNA donors as previously described (*69*). Modified cells were grown for 7-10 days then single cell sorted using a BD Aria II into 96 well U bottom plates. After 2-3 months of outgrowth, DNA from surviving clones (194/1200) was isolated using DNA quick extract solution (Lucigen) following a modified protocol. Briefly, 100 μL of cell culture was spun down, washed once with PBS, and then re-suspended in 20 μL of DNA extraction solution. Solution was heated in a thermocycler to 65°C for 15 minutes, 68°C for 15 minutes, 98°C for 10 minutes, and stored at 4 degrees. After DNA extraction, solution was diluted 1:20 and 5 μL used in a standard 50 μL PCR reaction using Q5 enzyme (NEB). PCR products were Sanger sequenced and analyzed using SnapGene to identify SNP corrected clones (7/192), wildtype HH clones (7/192), and a single (1/194) insertion / deletions clone. Sequences are shown in fig. S17.

### HDR sequence

ACAGCTCGGACCTGATGGATCTGATGTACCTGGCAGAAAGAATAAAAACCTGTGGATGTTTCCGTGAGTGGCAGGATTGGATGGTCGCTCGGAAAAGAACCAATCAGCACTGGAGCTGAAGGACCTC

### Real-time PCR and flow cytometry analysis of HH clones

For analysis of *HLA-DQB1* expression on HH WT (C/C) and ALT (G/G) clones, RNA was extracted from clones using a Monarch Total RNA extraction kit (NEB). cDNA was synthesized using MaximaH RT (NEB) enzyme following manufacturer’s protocol and oligoDT primers. cDNA was diluted 1 in 4 with *HLA-DQB1* (Assay ID:Hs00409790) and *actinB* (Assay ID:Hs01060665) probes and Taqman MasterMix (Thermofisher). Samples were run on an ARIAmx qPCR machine (Agilent) and data analyzed with Aria 1.5 (Agilent) software. Expression is represented as 2^−delta(*HLA-DQB*1 Ct - *ActinB* Ct)^. For analysis of protein expression, clones that survived after 3-4 months of culture were washed with PBS and stained with FITC anti-HLA-DQ (Biolegend, Clone: HLADQ1) for 30 minutes on ice and cell surface expression assessed by flow cytometry. Data was analyzed using Flowjo and Graphpad PRISM.

### Electrophoreitc Mobility Shift Assays

EMSAs were performed using the LightShift Chemiluminiscent EMSA Kit (Thermo Scientific). Single-stranded biotinylated oligonucleotides and complementary sequences corresponding to 31 nucleotides (15 nucleotides flanking the SNP of interest) were purchased from Eurofin Genomics and annealed by heating at 95°C for 5 minutes followed by a ramp down (−1°C/ minute) to room temperature (20°C).

Nuclear extract from Jurkat, HH, and Daudi cells was isolated using the NE-PER nuclear and cytoplasmic extraction kit (Thermofisher Scientific) with slight modification. 20 million cells were spun down, washed twice with PBS, and used in the protocol with half volumes of NER buffer. All buffers contained protease inhibitors (Thermofisher). Protein extracts were then dialyzed using a membrane with a molecular weight cut-off of 12–14 kDa (Spectrum Spectra) against 1 L of dialysis buffer (10 mM Tris pH 7.5, 50 mM KCl, 200 mM NaCl, 1 mM dithiothreitol, 1 mM phenylmethane sulfonyl fluoride, and 10% glycerol) for 16 h at 4°C with slow stirring. The protein concentration was measured using the Pierce BCA protein assay kit (Thermo Scientific).

The standard binding reaction contained 2 μL of 10× Binding Buffer, 2.5% glycerol, 5 mM MgCl2, 0.05% NP40, 50 ng Poly(dI:dC), 20 fmol biotin-labeled probe, and 10-20 μg of nuclear extract in a final volume of 20 μL. For competition, a 200-fold molar excess (4 pmol) of unlabeled probe was added.

Binding reactions were incubated at room temperature for 30 min and loaded onto a 6% polyacrylamide 0.5 × Criterion precast TBE gel (Biorad). After sample electrophoresis and transfer to a nylon membrane, DNA was crosslinked for 10 min, and biotinylated probes detected by chemiluminescence followed by film exposure. Original films and replicated are presented in a fig. S19.

List of probes:

Alt A CAGGATTGGATGGTCGCTCGGAAAAGAACCA
Alt B TGGTTCTTTTCCGAGCGACCATCCAATCCTG
Ref A CAGGATTGGATGGTCCCTCGGAAAAGAACCA
Ref B TGGTTCTTTTCCGAGGGACCATCCAATCCTG

### Luciferase assay

A double-stranded oligonucleotide containing the SNP of interest (31nt + restriction enzyme sites) was ordered and annealed as described above. Probes and the luciferase reporter vector pGL3 promoter (Promega) were digested with BglII (NEB) for 1 h at 37°C, and the linearized vector was simultaneously dephosphorylated with alkaline phosphatase (NEB). Digestion products were purified with the Gel Extraction Kit (Thermofisher) from 1% agarose gels. Ligation was then performed in a ratio of 1:50 (vector:insert) with T4 DNA ligase (NEB) at room temperature for one hour and then transformed into NEB10 competent cells (NEB). Plasmids from independent colonies were isolated using a plasmid DNA minikit and Sanger sequenced to identify correctly inserted clones.

3 × 10^5^ Jurkat, HH, and Daudi cells were nucleofected with 0.9 μg of pGL3-Promoter vector along with 0.1 μg of pRL-TK Renilla luciferase vector (Promega) in 16 well strips in a 4D nucleofector with the following protocols and buffers in 20 μL of total volume: Jurkat, SE buffer, CL-120 protocol; HH, SE buffer, CL-120 protocol; Daudi SF buffer, CA-137 protocol. After nucleofection, 180 μL of complete RPMI was added and cells cultured in 96 well flat bottom plates (Falcon). After 48 hours, cells were spun down, resuspended in 75 of fresh complete media, and luciferase/renilla activity measured using the Dual-Glo Luciferase Assay System (Promega). Firefly luciferase activity was expressed as relative luciferase units (RLU) after correction for Renilla luciferase activity. Data were normalized to those cells transfected with empty pGL3-Promoter vector. Each dot represents an independent nucleofection reaction.

### Autoimmune disease enrichment analyses

We downloaded SNPs from the GWAS catalogue on July 17, 2018. We selected SNPs with P < 5e-08 for 11 autoimmune diseases and 3 non-immune mediated diseases that served as a negative control (schizophrenia, type 2 diabetes, coronary heart disease). We used SNPsea to capture genes within disease loci based on LD and recombination interval information (*70*). We then assessed how many of the dynASE genes overlap genes in disease loci for each disease (observed overlap). To assess whether this overlap represented a significant enrichment, for each disease we created 1000 null sets of N random regions in the genome (N = number of disease loci), which were matched by the number of genes per locus (within 15% of each disease locus). We then calculated the ratio of observed overlap with the mean overlap of our 1000 null sets (fold enrichment). We calculated the P-value as: (number of null overlaps larger than observed overlap + 1)/1001 (*71*). We took the same approach for genes with significant ASE at 0 hours (N = 501), and eQTL genes of naïve T cells reported by the Blueprint Consortium (N = 5,688) (*8*).

We carried out a heritability enrichment analysis using GWAS summary statistics and stratified LD score regression (S-LDSC) (*32*) for six autoimmune diseases and three non-immune mediated diseases. For each gene category described above (dynASE, 0 hours ASE, naive T eQTL), we separately created functional annotations consisting of all SNPs within +/− 100 kb of each gene. We define enrichment of an annotation as the proportion of heritability explained by the annotation divided by the proportion of the common (MAF ≥ 0.05) SNPs included in the annotation. To test for enrichment significance, S-LDSC tests if the per-SNP heritability is greater in the functional annotation than outside of the functional annotation. Standard errors around the enrichment mean are computed by using a block jackknife of 200 equally-sized adjacent blocks of SNPs genome-wide. Then, we compute a z-score to test for significance.

## Acknowledgments

We are indebted to Gila Klein RN for her outstanding management of the Genotype and Phenotype (GaP) registry at the Feinstein Institute, to the Raychaudhuri laboratory members for critical discussions and feedback, and to Henry Long and Paloma Cejas for support on primary T cell ATAC-seq experiments.

## Author Contributions

M.G.-A., Y.B., and S.R. conceived, designed, and performed analyses/experiments, wrote the paper, and supervised the research. J.A., S.H., Y.L., T.A., N.T. performed analyses or experiments, interpreted data, and contributed to the paper. D.A.R., J.E., A.H.J., M.B.B. provided experimental supervision and contributed to the paper. C.N., P.K.G., contributed to sample recruitment and results discussion, C.N. facilitated experiments, T.E. contributed to data acquisition.

## Data and materials availability

All data created for this study will be made available in GEO and dbGAP.

---

**Fig. S1.**
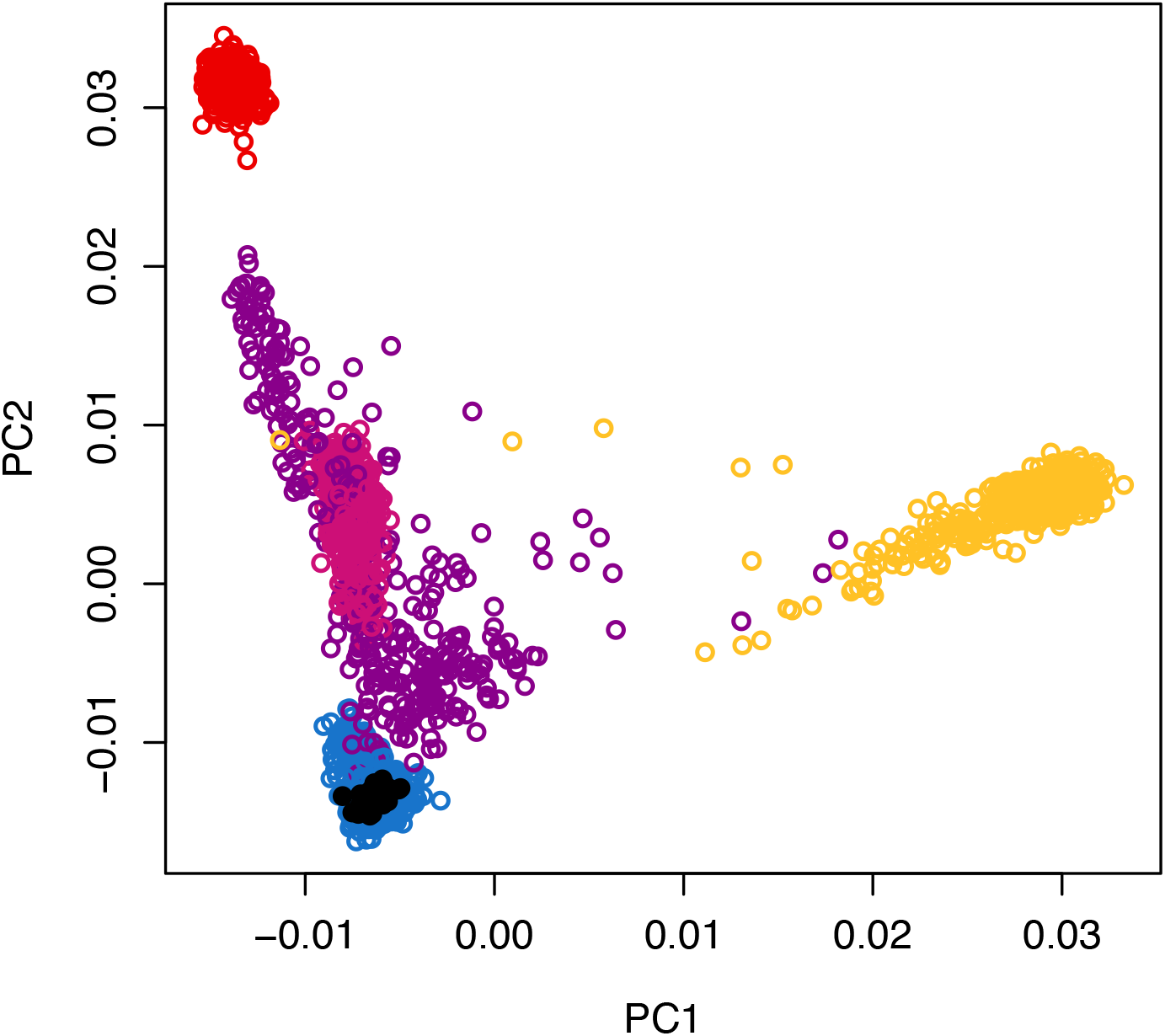
Individuals of European ancestry. Principal component analysis performed on 8,772 LD pruned SNPs with MAF of 5% from 2,238 individuals from the GaP Registry, and 2,504 individuals from the 1,000 genomes project. Shown are only the 24 GaP individuals that were recruited for the present RNA-seq study (black), and all the 1,000 genome individuals of European (blue), African (yellow), East Asian (red), South Asian (pink) and Ad Mixed American (purple) ancestries.

**Fig. S2.**
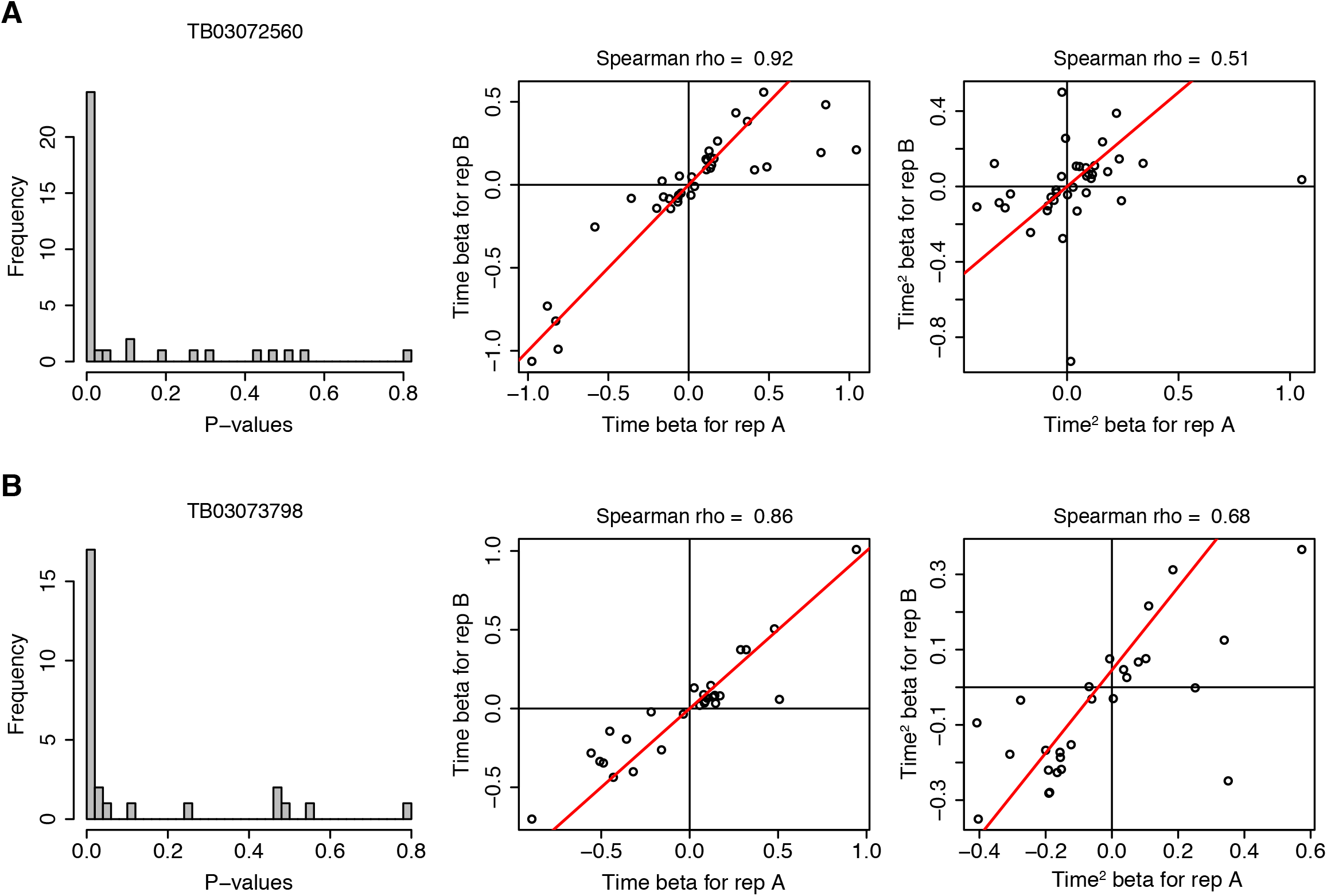
Replication of dynamic ASE in two pilot individuals. **(A)** For dynamic ASE events significant in replicate A of individual TB03072560, the P-values (left panel) and beta for time (middle) and time squared (right) were checked in replicate B. **(B)** Same as for (A) but for individual TB03073798.

**Fig. S3.**
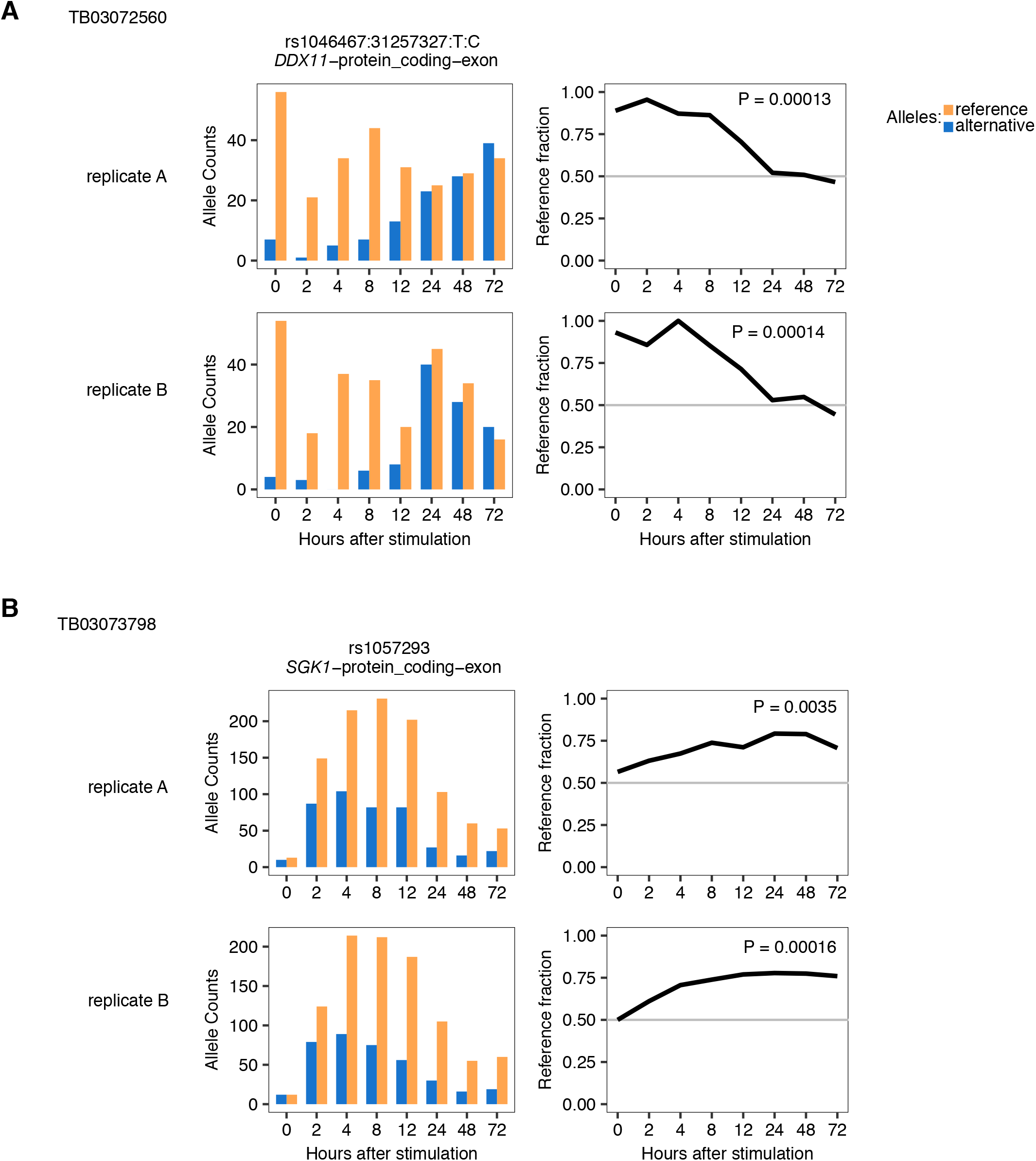
Replication examples of dynamic ASE in two pilot individuals. Examples of dynamic ASE events significant in replicate A of individual TB03072560 **(A)**, and replicate A of individual TB03073798 **(B)**. Shown are allelic counts for heterozygous SNP (left) and reference fraction over time (right) for replicate A (top panels) and replicate B (bottom panels).

**Fig. S4.**
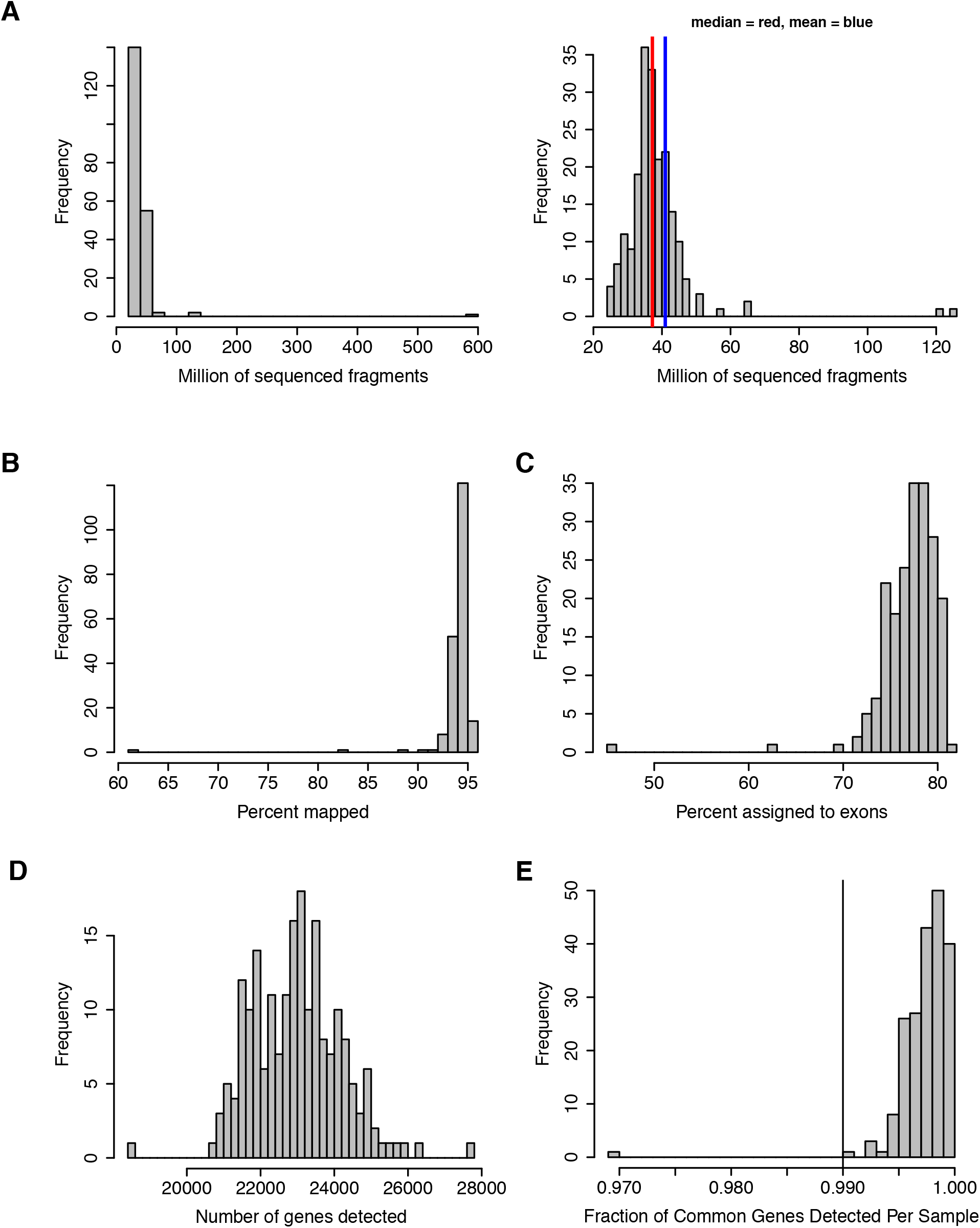
RNA-seq data summary and quality check. **(A)** Histograms showing the number of sequenced fragments (read pairs) per library. Right panel is a zoom in of left panel, with median (red, 37M) and mean (blue, 41M) indicated. **(B-D)** Histograms showing percent of mapped fragments **(B)**, percent of fragments assigned to exons **(C)** and number of genes detected (>=1 mapped fragment, **D)** per library. **(E)** Histogram showing the fraction of common genes detected per sample. Common genes defined by being detected in >95% of samples.

**Fig. S5.**
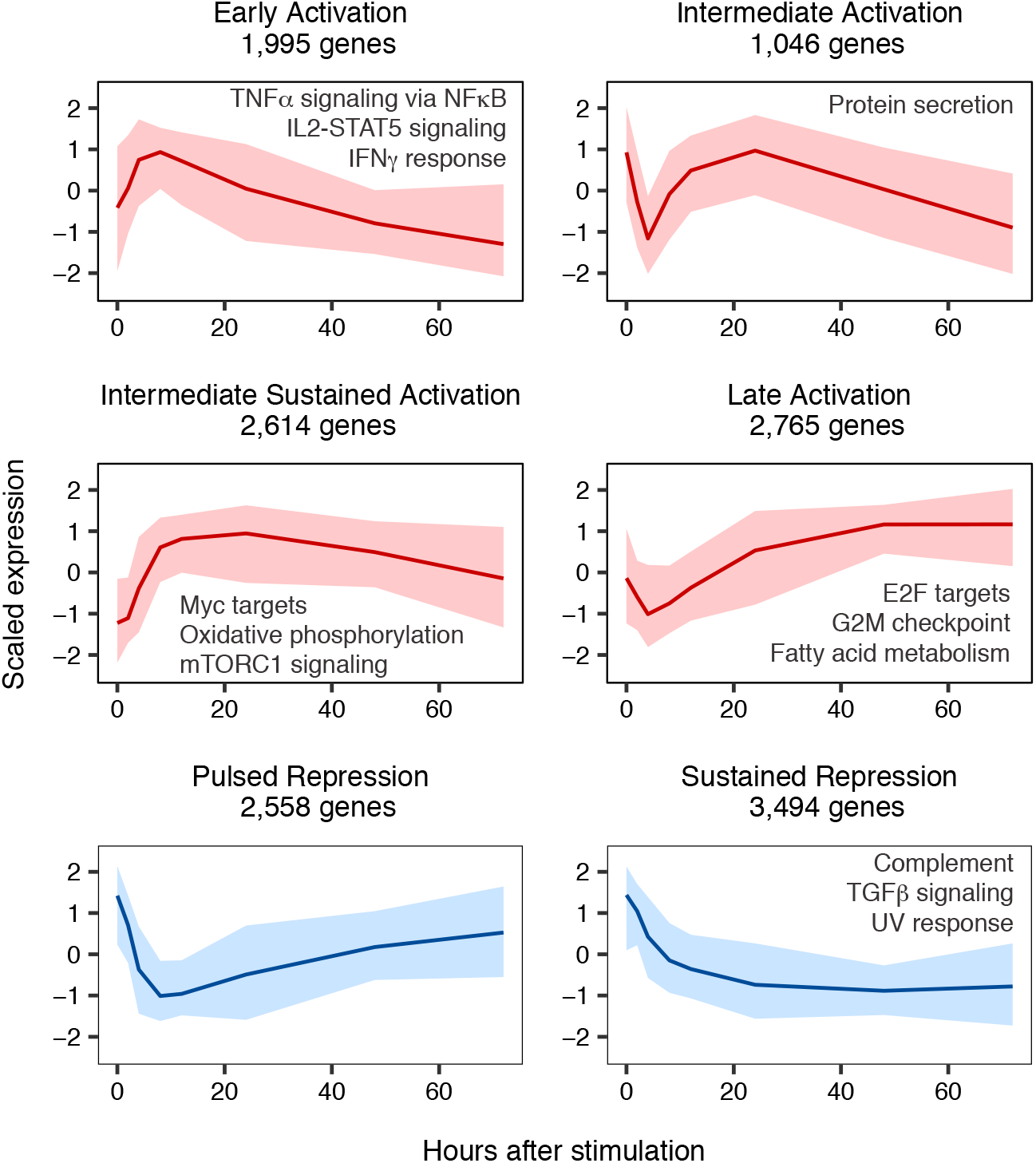
Six gene expression clusters show activated and repressed genes and pathways. Scaled log2(tpm+1) gene expression levels for genes pertaining to each of the four activation clusters (red) and two repression clusters (blue) detected through k-means clustering. Central line indicates mean expression per cluster. Ribbons around the central line show the 2.5 and 97.5 percentiles of expression among all genes in the cluster. Indicated in each plot are pathways significantly enriched for genes in the cluster.

**Fig. S6.**
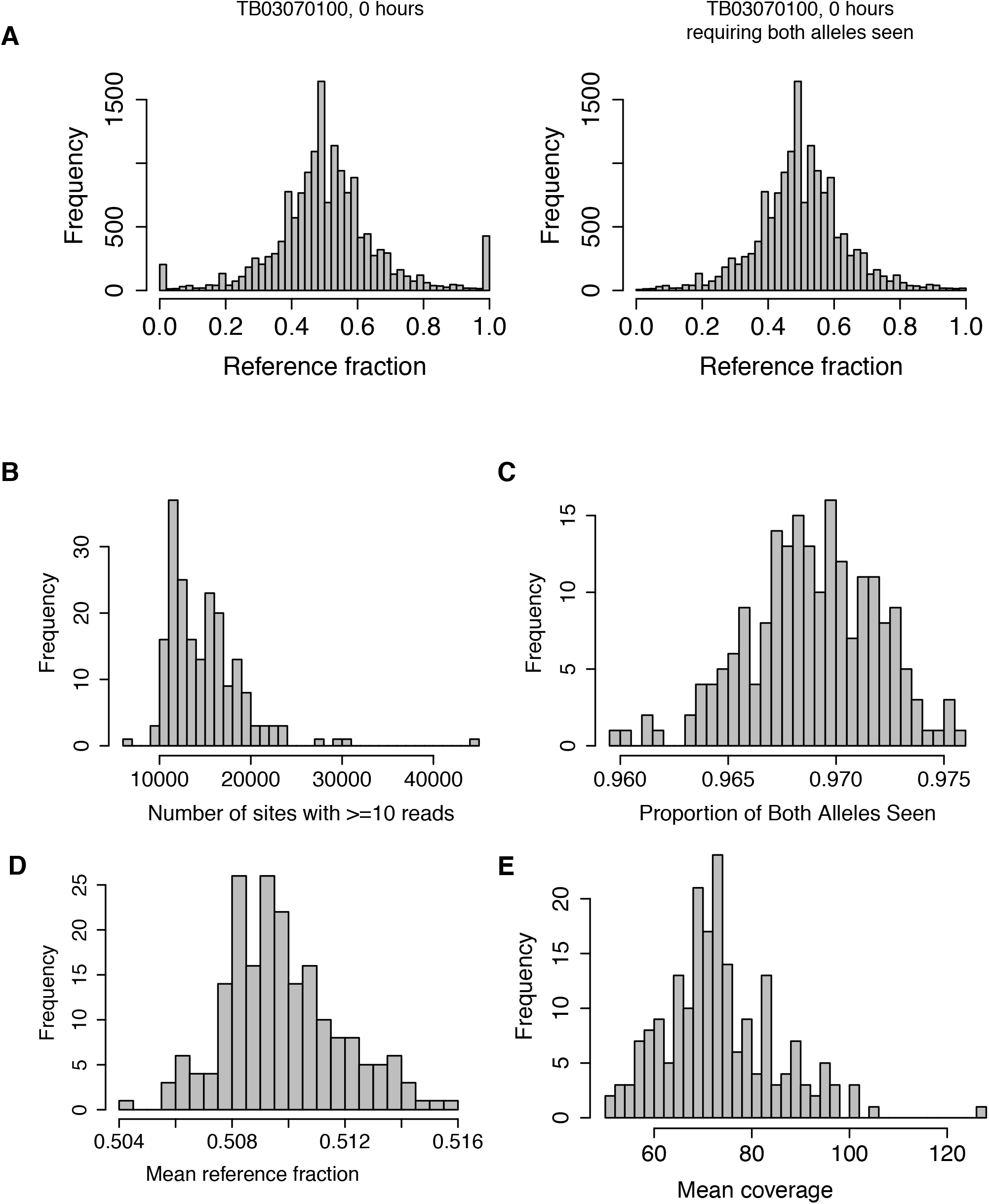
Allele specific expression data summary and quality check. **(A)** Histograms showing the reference fraction distribution of a typical sample. Included are all heterozygous sites with at least 10 reads (left), and requiring both alleles seen (right). Histograms showing per sample, the number of sites with at least 10 reads **(B)**, and then within, sites with at least 10 reads: the proportion of both alleles seen **(C)**, the mean reference fraction **(D)** and mean coverage **(E)**.

**Fig. S7.**
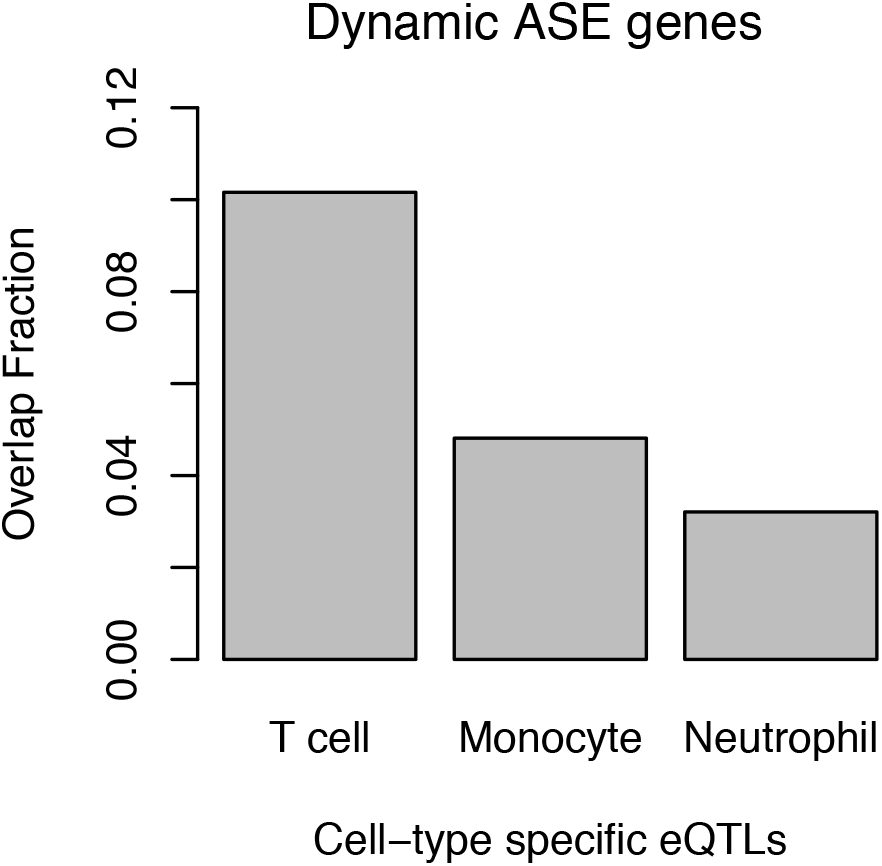
Dynamic ASE genes are enriched for T cell specific eQTL genes. Overlapping fraction of dynamic ASE genes (5% FDR) with Blueprint-reported eQTL genes identified in naive T cells but not monocyte or neutrophils (T cell), in monocytes but not T cells or neutrophils (Monocyte), and in neutrophils but not T cells or monocytes (Neutrophil).

**Fig. S8.**
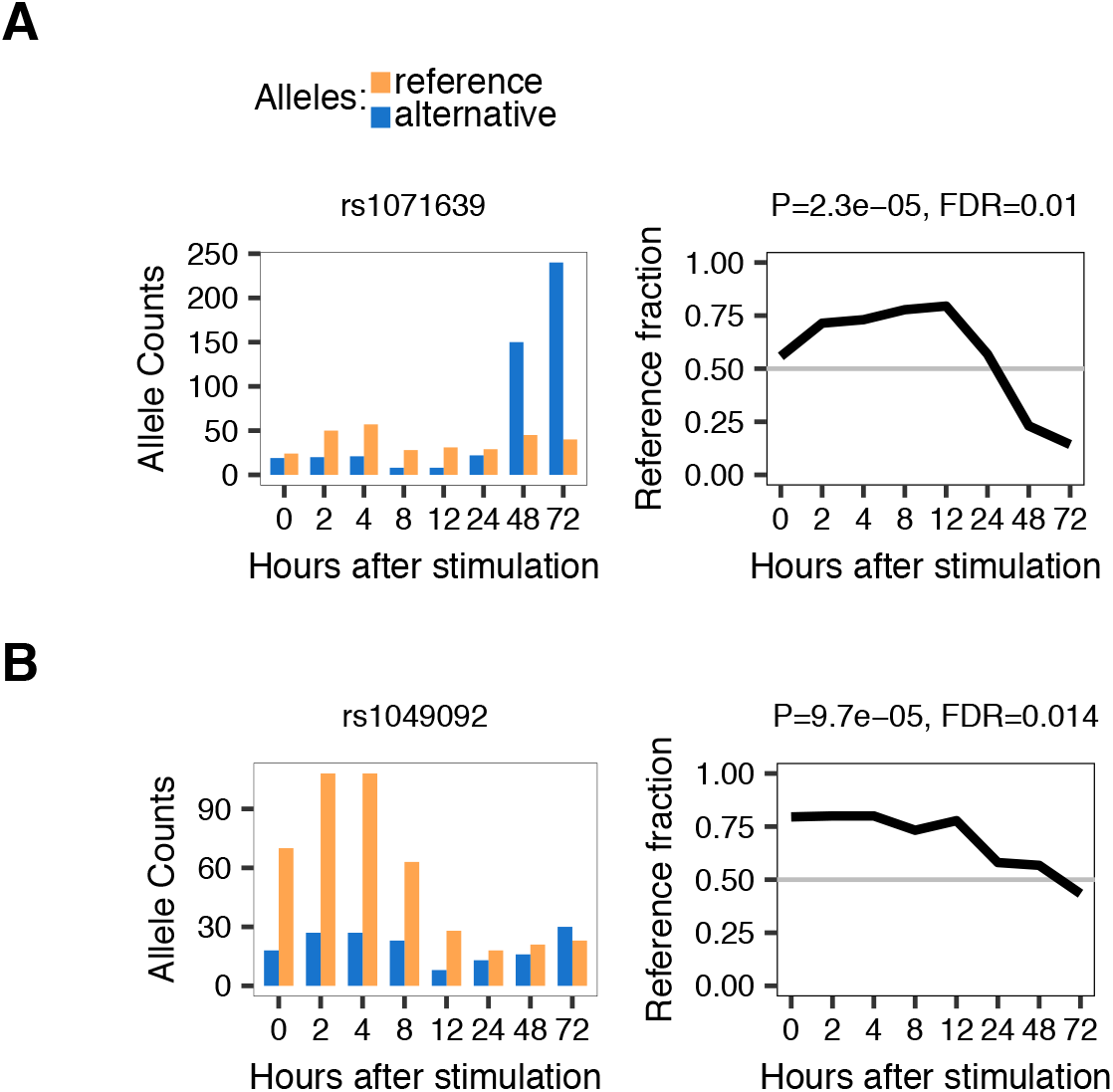
Dynamic ASE in *HLA-DQB1* SNPs. Allele counts (left panel) and the correspoinding reference fraction over time (right panel) for two heterozygous SNPs in *HLA-DQB1* for individual TB03074401 **(A)** and TB03073148 **(B)**.

**Fig. S9.**
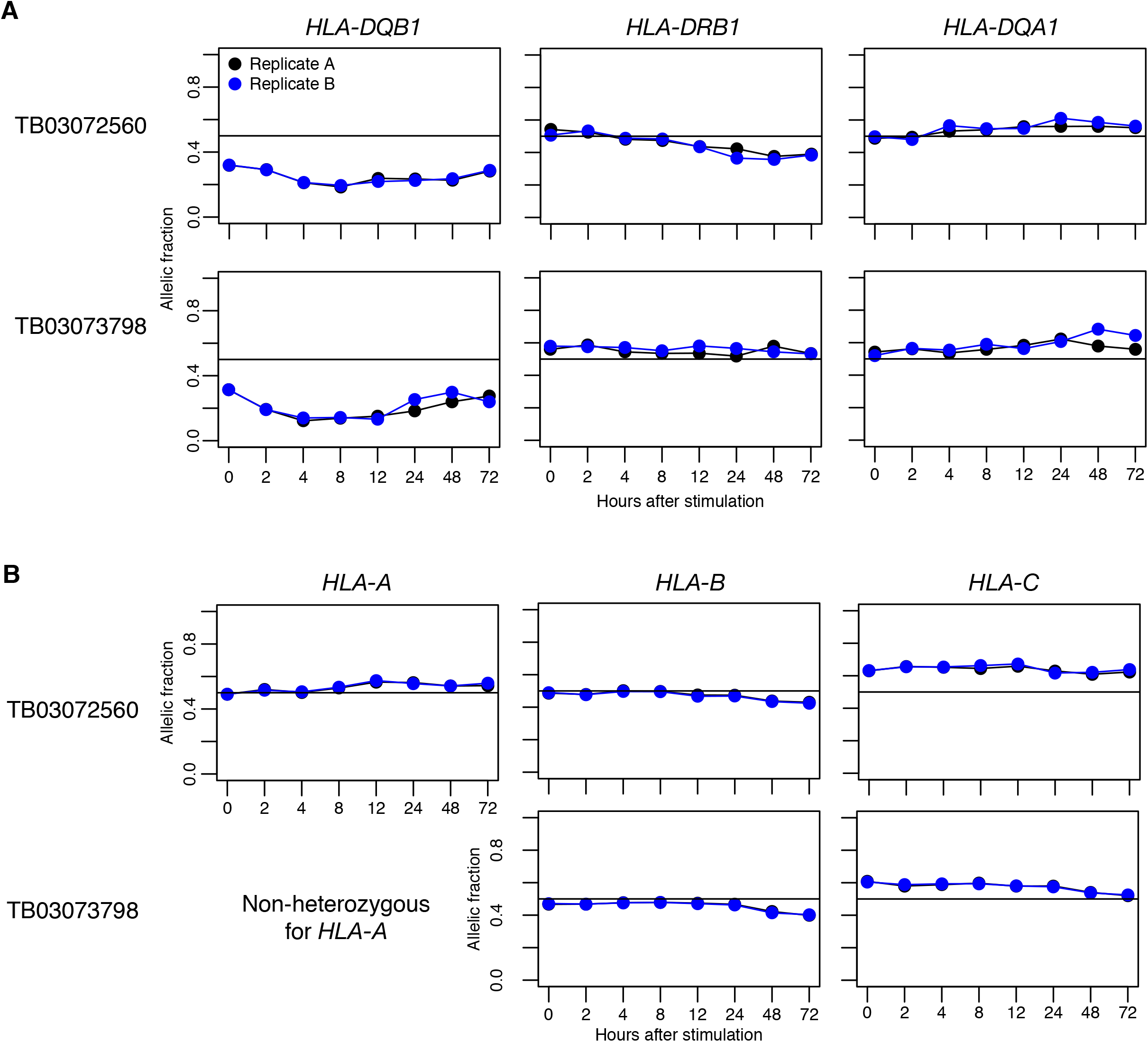
Allelic fraction replication in HLA gene quantifications. Allelic fraction across time for the 3 HLA class II genes **(A)** and 3 HLA class I genes **(B)**, for the two pilot individuals with full time course replicates. Replicate A in black, replicate B in blue.

**Fig. S10.**
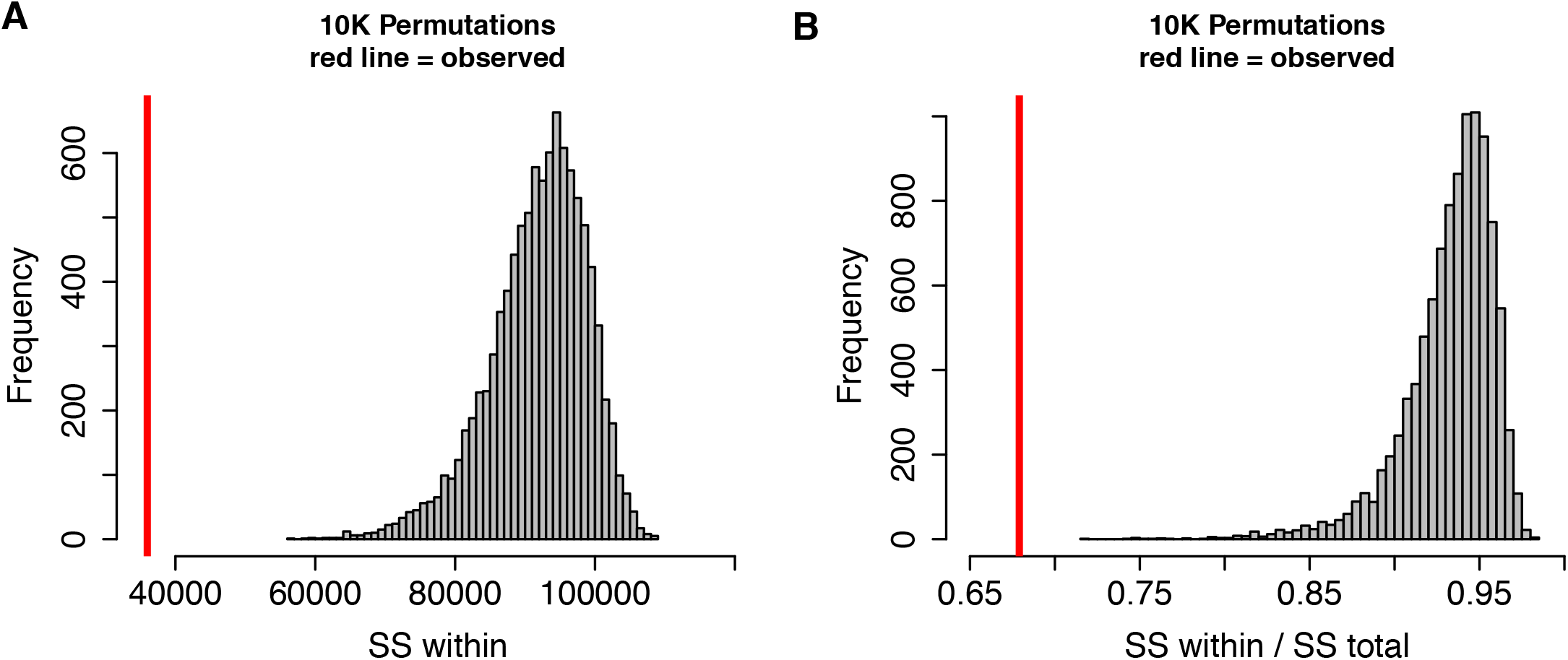
*HLA-DQB1* 4-digit allele groups have similar allelic profiles among each other. Histograms showing distribution of **(A)** sum of squares within 4-digit classical allele groups, and **(B)** sum of squares within 4-digit classical allele groups over total sum of squares, for 10,000 permutations of 4-digit classical alleles on *HLA-DQB1* allelic expression profiles. Red line indicates observed value.

**Fig. S11.**
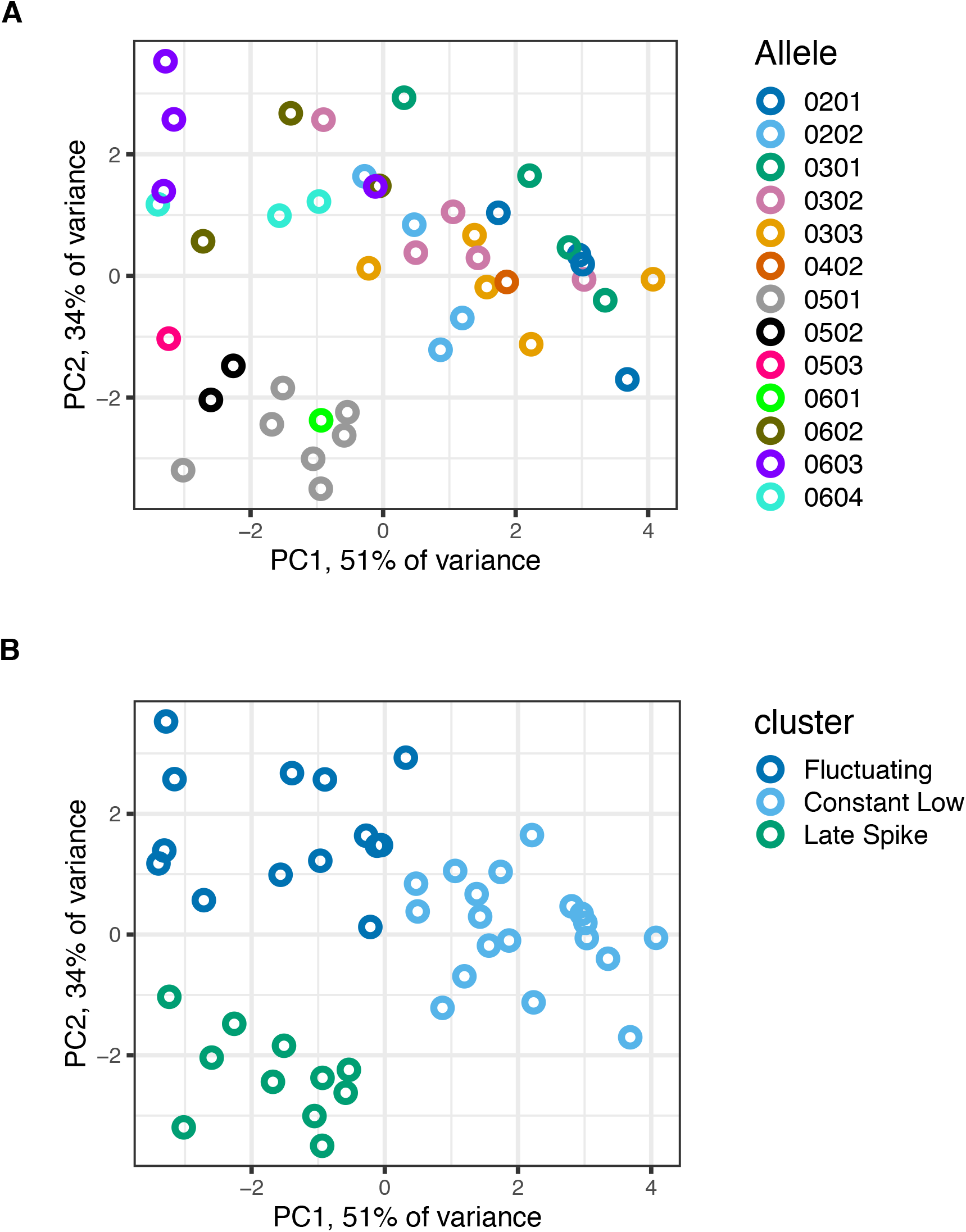
Pincipal component analysis of *HLA-DQB1* allelic profiles over time. PCA performed for 48 *HLA-DQB1* allelic profiles of 24 individuals (log2(FPKM+1) values over time, colored by 4-digit classical *HLA-DQB1* allele **(A)**, and by the k-means cluster to which they belong **(B)**. Average allelic expression was computed for samples with replicates. Twelve hour time point was removed because of high number of missing values.

**Fig. S12.**
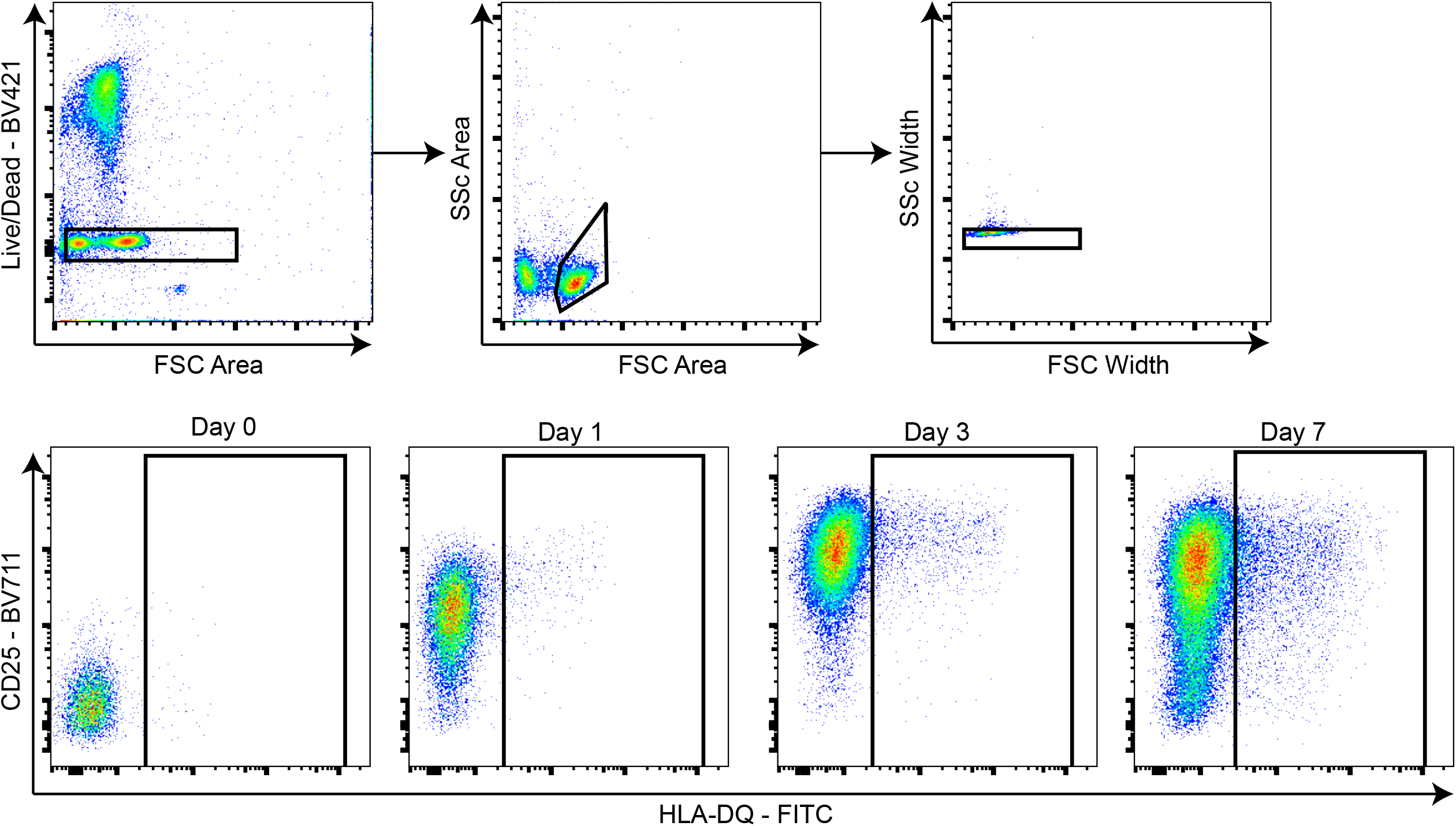
Representative flow cytometry staining of stimulated T cells from PBMCs. Memory CD4^+^ T cells from a healthy donor homozygous for a *Late-Spike* allele stimulated with anti-CD3/CD28 microbeads for 7 days, and gated as shown to measure median flourescence intensity of HLA-DQ.

**Fig. S13.**
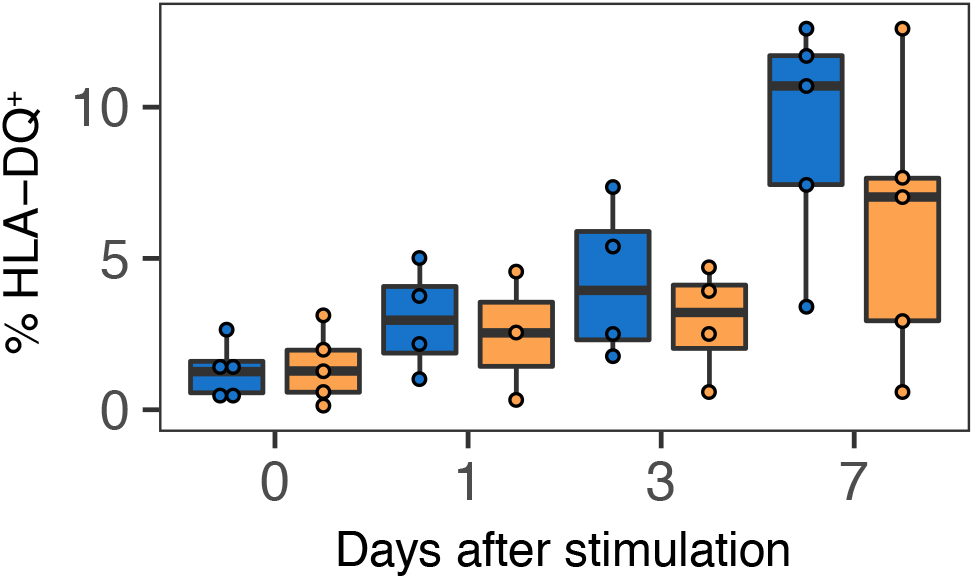
Percent HLA-DQ+ cells within CD4^+^ memory T cells over time. In blue, individuals homozygous for *Late-Spike* alleles. In yellow, individuals homozygous for *Constant-Low* or *Fluctuating* alleles.

**Fig. S14.**
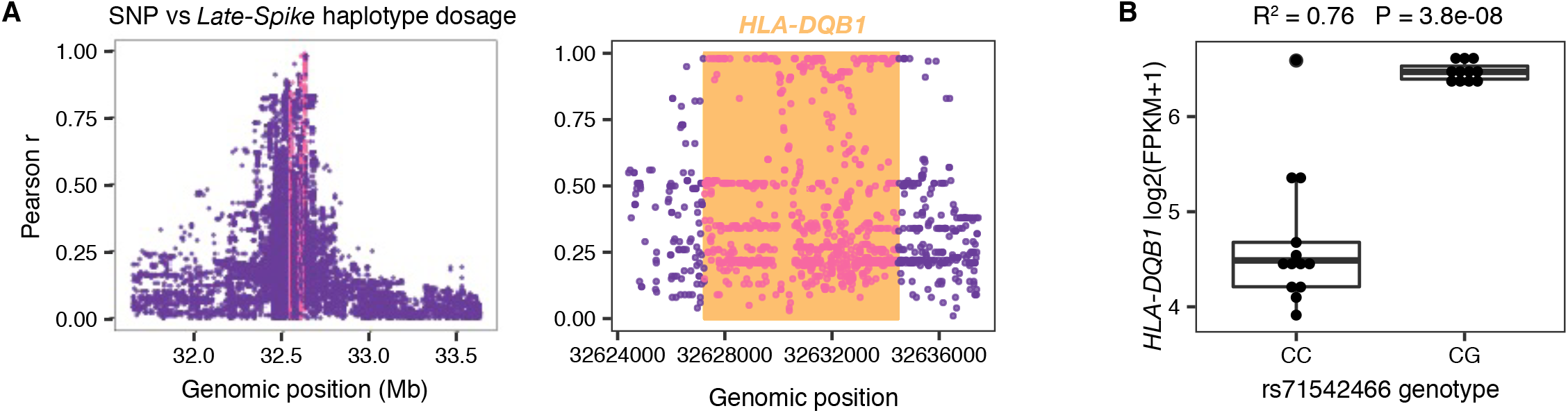
Mapping variants associated with Late Spike haplotype. **(A)** Pearson correlation coefficient between SNP genotypes and *Late-Spike* haplotype dosage. Orange vertical lines indicate location of *HLA-DRB1, HLA-DQB1* and *HLA-DQA1* genes, where dots are colored pink. Right plot is zoomed in on *HLA-DQB1* region to show top SNPs. **(B)** *HLA-DQB1* gene expression levels (log2(FPKM+1)) for individuals separated by their rs71542466 genotype.

**Fig. S15.**
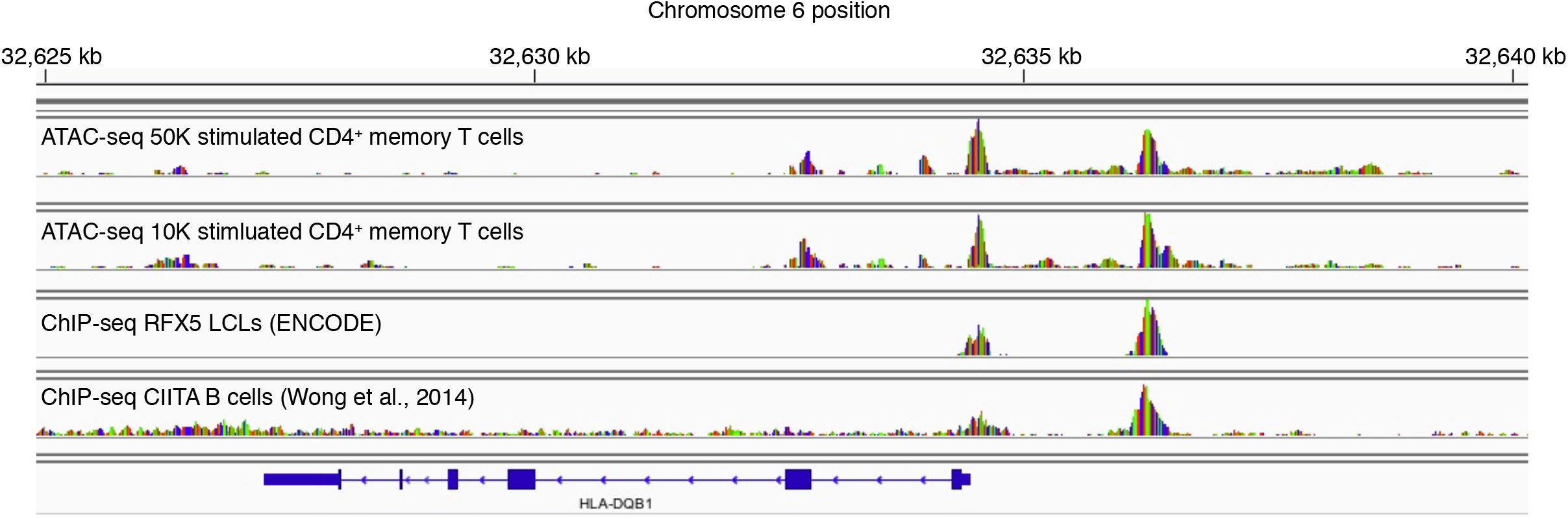
Epigenomic landscape around *HLA-DQB1*. IGV read pileup tracks for ATAC-seq performed on 50K or 10K cells on CD4^+^ memory T cells stimulated for 72 hours with anti-CD3/CD28 microbeads, ChIP-seq from published data for RFX5 in LCLs, and for CIITA in B cells.

**Fig. S16.**
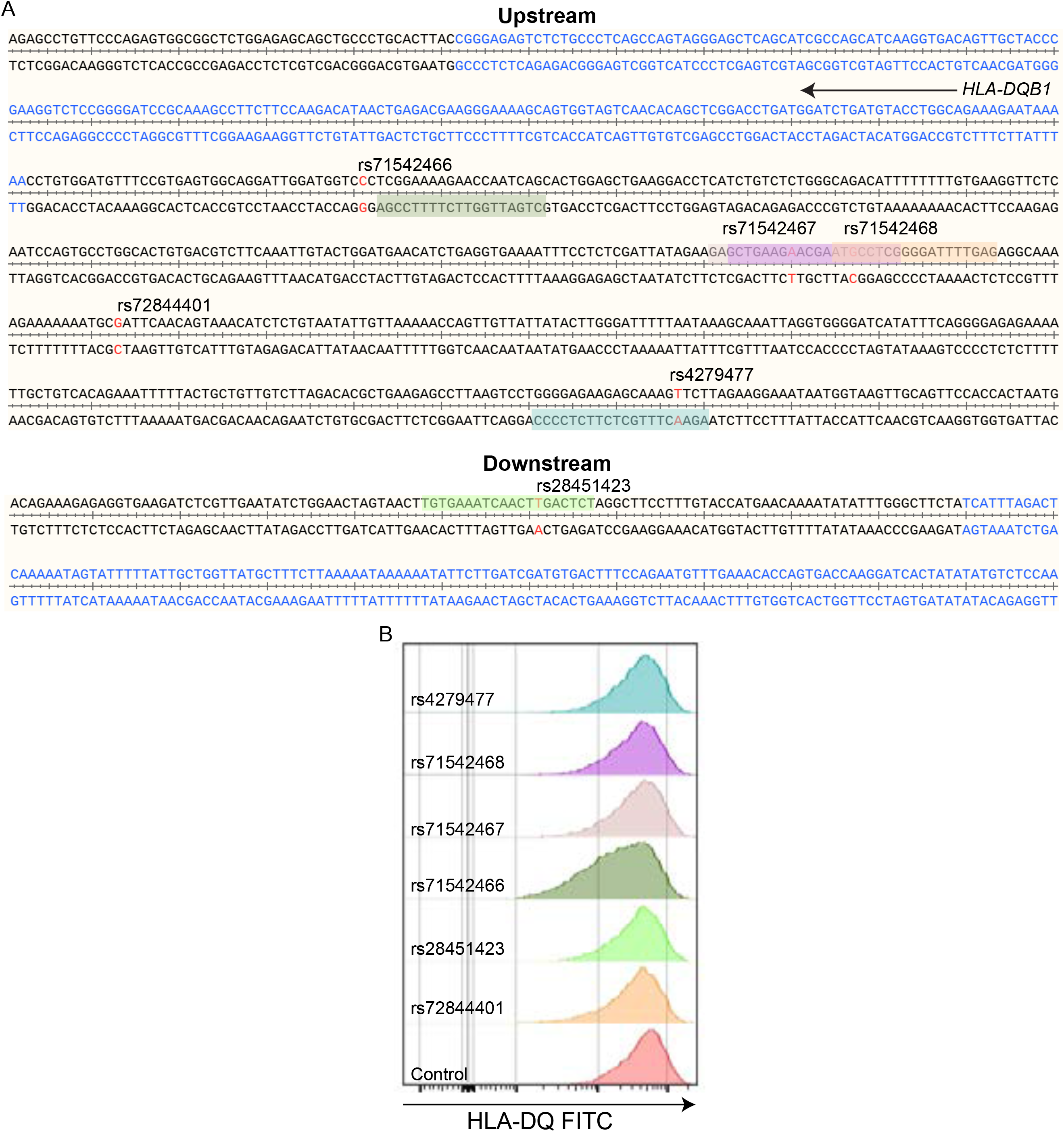
Genomic location of nearest gRNAs to identified SNPs and representative flow cytometry plot of CRISPR-Cas9 edited HH cells. **(A)** Location of SNPs (red colored nucleotides) is shown in reference to the nearest exon (blue colored nucleotide) both upstream and downstream of *HLA-DQB1*. The nearest gRNA sequences are highlighted with their corresponding colors (rs71542466 - dark green, rs71542467 - light purple, rs71542468 - purple, rs72844401 - beige/orange, rs4279477 - blue, rs28451423 - light green). Alignments were plotted using SnapGene(v3.2.1). **(B)** Representative staining of HLA-DQ of modified HH cells with corresponding CRISPR-Cas9 complexes. Cells were stained 7-10 days after modification as a bulk population.

**Fig. S17.**
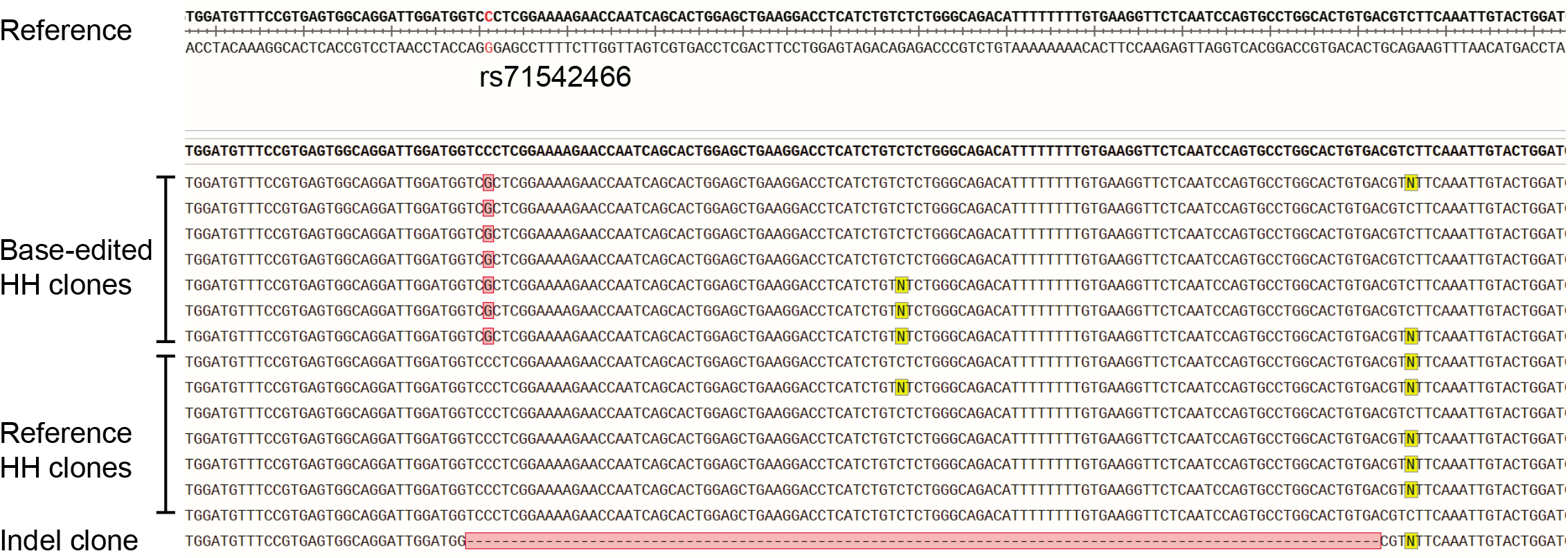
Sanger sequencing alignment of HH reference and base-edited clones reveal seamless editing. gDNA from expanded clones was sequenced and aligned to the reference (hg38) and visualized using SnapGene(v3.2.1). Red colored nucleotide indicates the location of the rs71542466 SNP. Highlighted red nucleotides indicate mismatches from the reference and yellow coloured nucleotides indicate unresolved/heterozygous sequences.

**Fig. S18.**
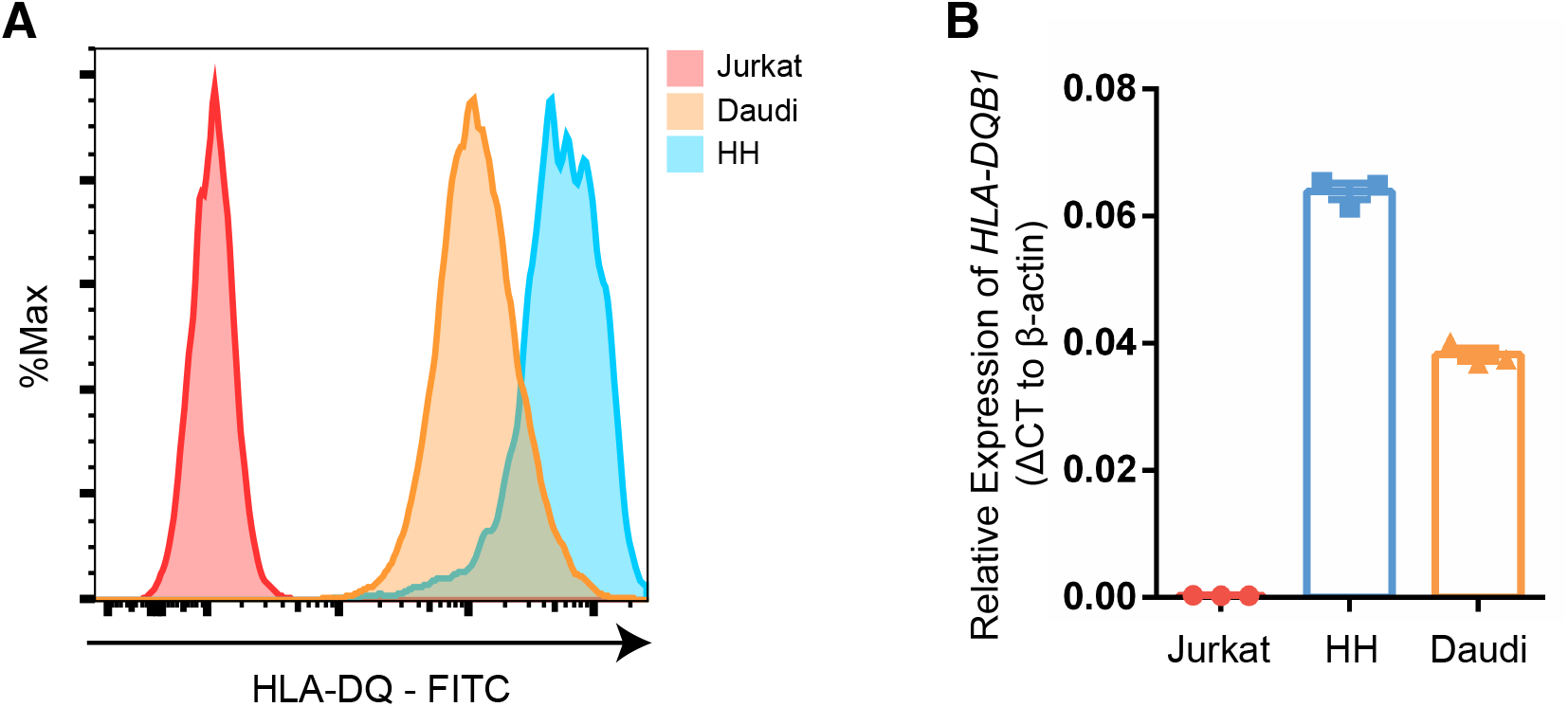
Expression of *HLA-DQB1* on Jurkat, HH and Daudi cells. **(A)** Histograms showing the cell surface expression of HLA-DQ as measured by flow cytometry. **(B)** Expression of *HLA-DQB1* as measured by real-time PCR. Each dot represents an independent RNA extraction (N = 3).

**Fig. S19.**
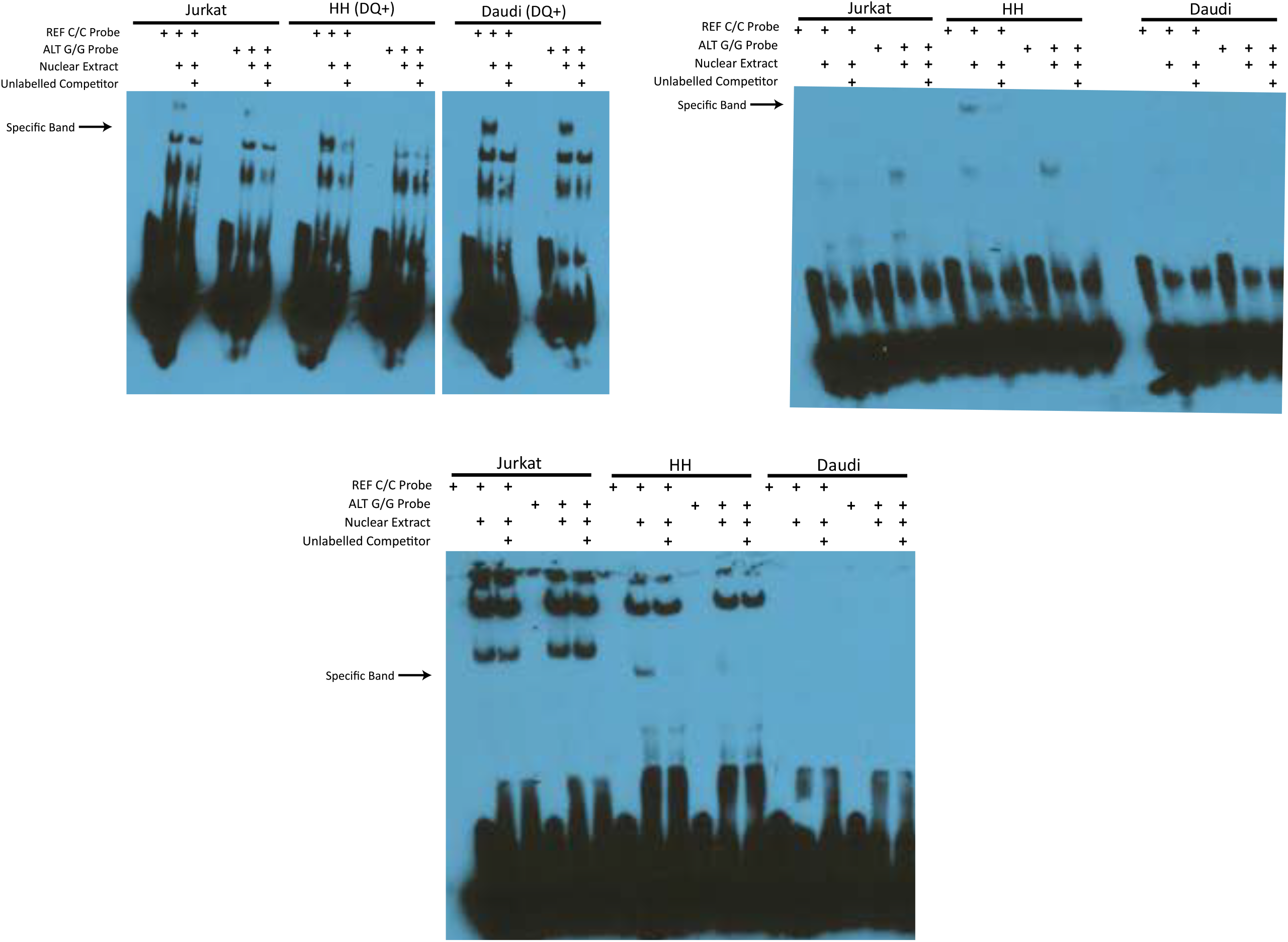
Independent EMSAs identifying a specific binding band in HH cells. EMSA asays were performed on three independently isolated nuclear extractions to validate the presence of a specific nuclear binding band in the REF allele. 20pmol of biotinylated probe was mixed with 10-15ug of nuclear extract in each independent replicate.

**Fig. S20.**
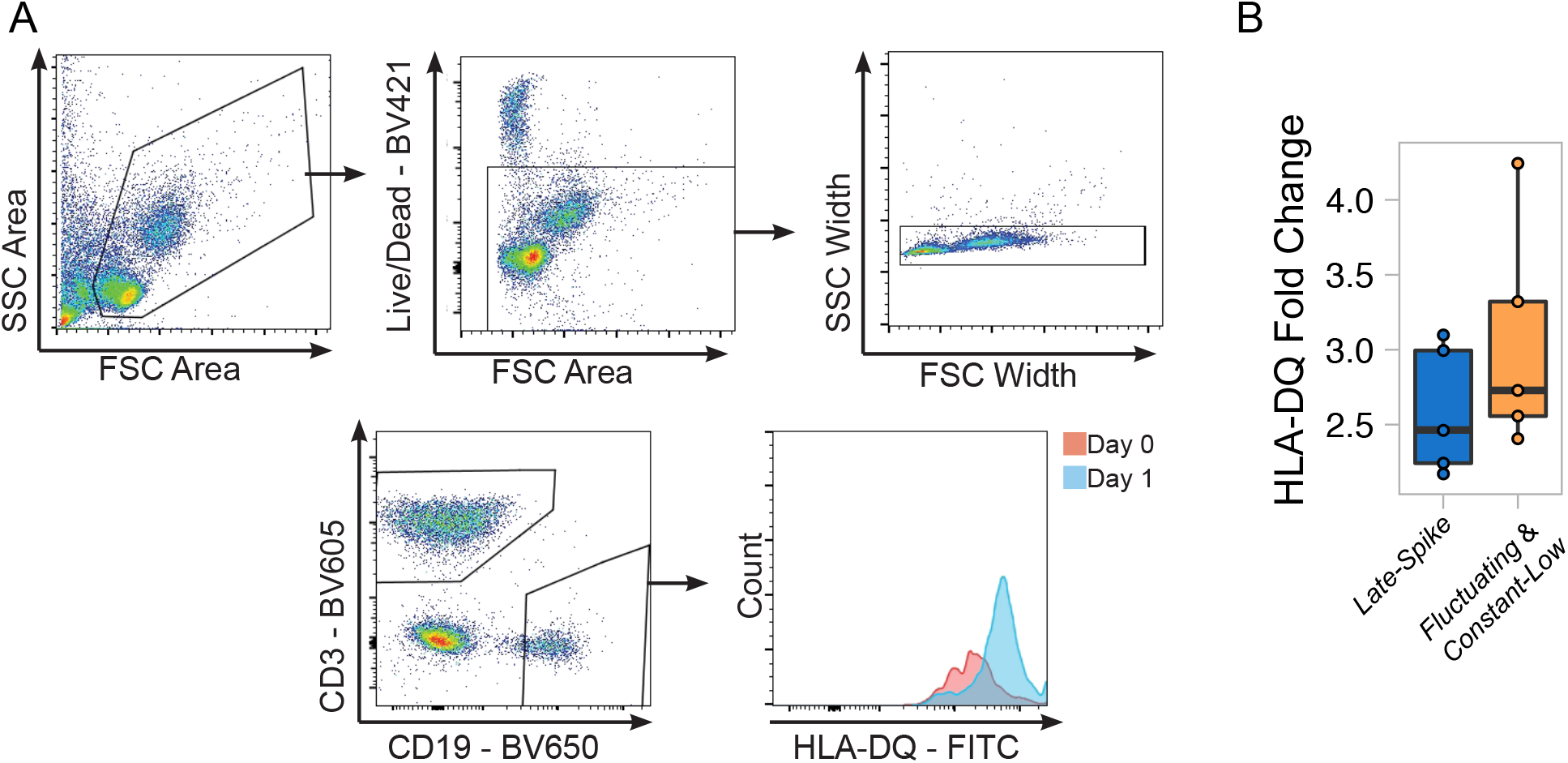
Representative flow cytometry staining of stimulated B cells from PBMCs and fold change of HLA-DQ. **(A)** Representative plot of stimulated B cells from *Late-Spike* control PBMCs. Cells were stimulated with anti-Ig, CD40L, rhIL-21, and CpG(2006) for 1 day. **(B)** Summary of median flourescence intensity fold change in HLA-DQ expression from day 0. Each dot represents an individual.

**Fig. S21.**
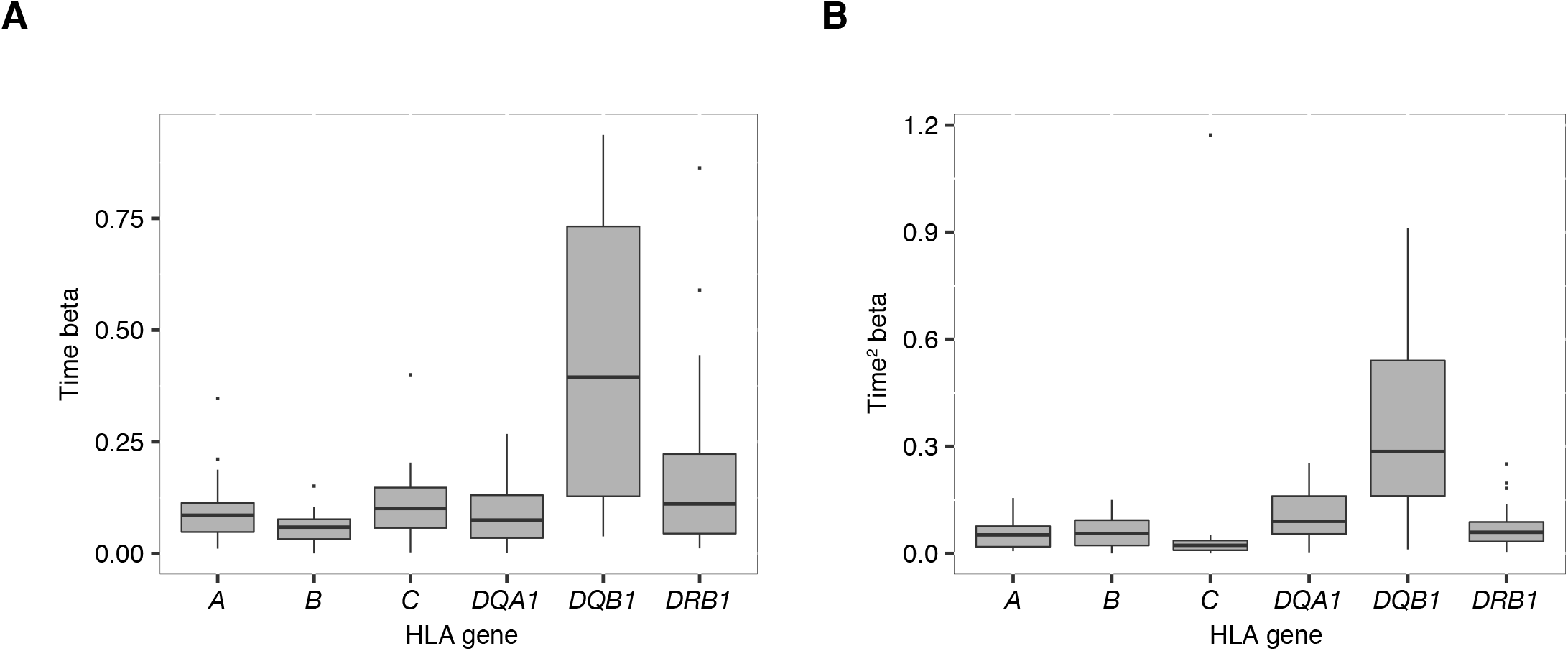
Dynamic ASE betas in HLA genes. Distributions of beta for time **(A)** and time squared **(B)** per HLA gene.

**Fig. S22.**
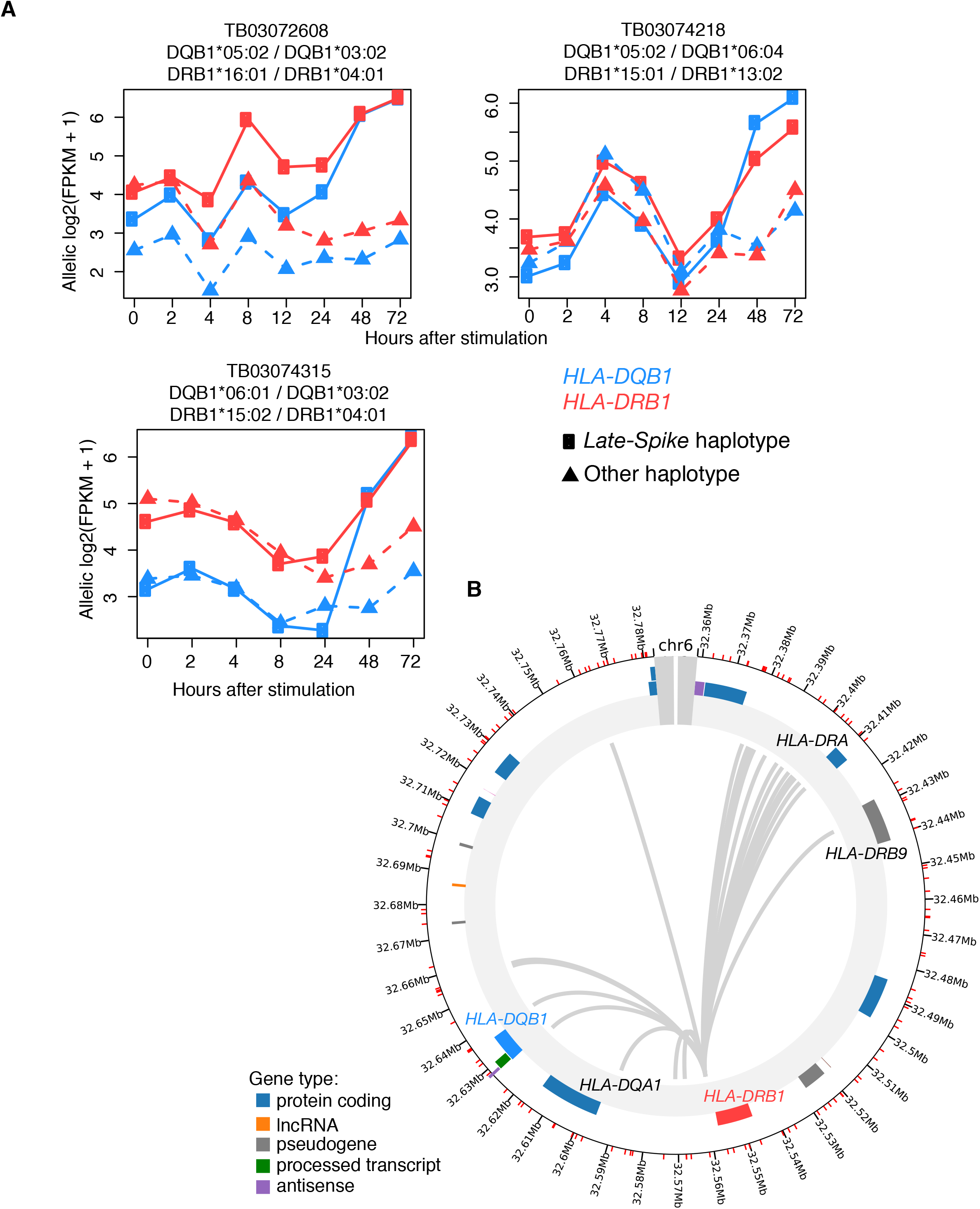
*HLA-DQB1* and *HLA-DRB1* haplotypes and chromosomal interactions. **(A)** Three individuals with *HLA-DQB1 Late-Spike* haplotype that have an *HLA-DRB1 Late-Spike-like* allelic expression profile. Alleles on one chromosome indicated on the left (*Late-Spike* haplotype), and alleles on the other chromosome indicated on the right (other haplotype) on plot titles. **(B)** Promoter capture HiC interactions for *HLA-DRB1* (red). Image adapted from www.chicp.org, data comes from Javierre et al Cell 2016.

**Fig. S23.**
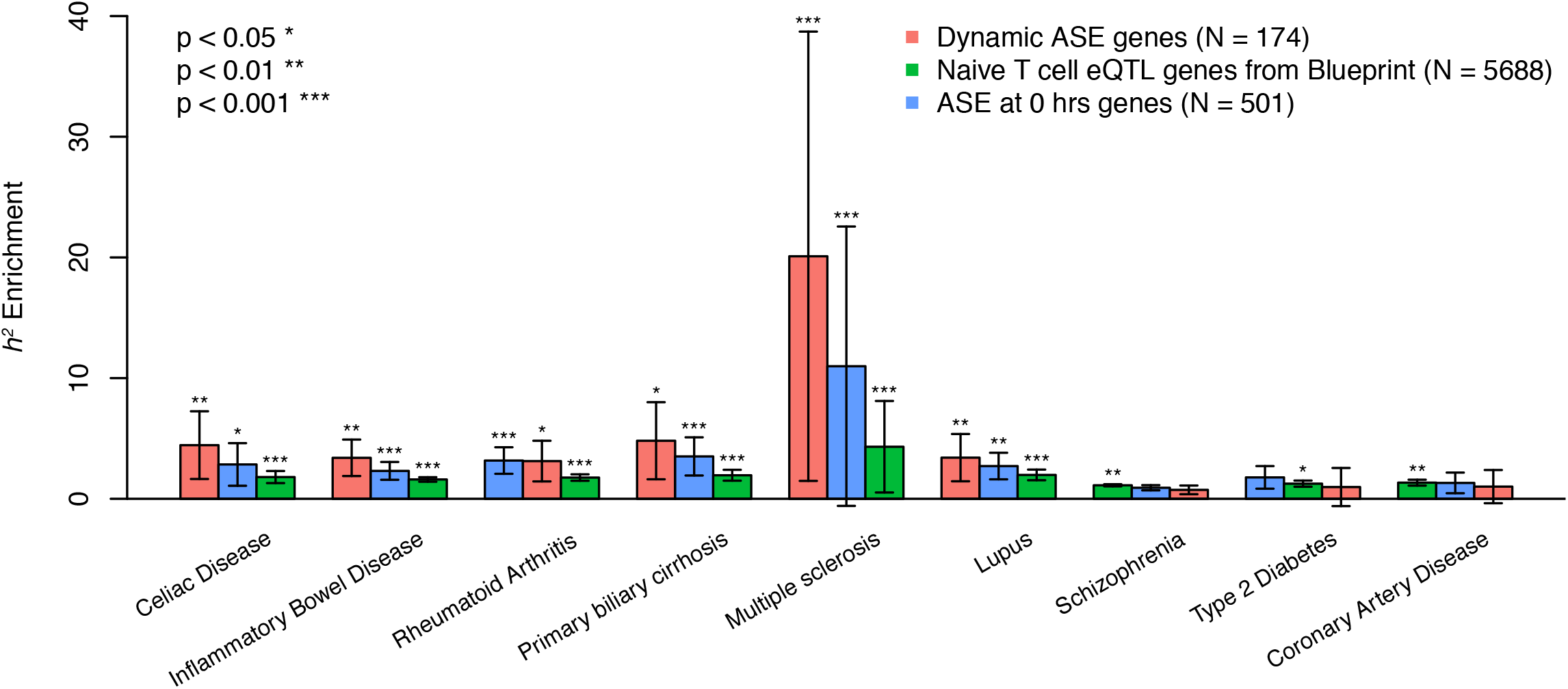
Disease heritability enrichment for genes with different types of regulatory effects. Heritability enrichment calculated with GWAS summary statistics and stratified LD score regression. Gene annotations include gene +/− 100kb.

**Table S2.** Reported eQTLs for *HLA-DQB1* and LD with the *Late-Spike* regulatory SNP

| Cell-type and condition | Study | Read mapping method | SNP ID | SNP chr6 position | SNP position relative to TSS | r2 with rSNP |
| --- | --- | --- | --- | --- | --- | --- |
| 48-72 hrs stimulated CD4+ memory T cells | present | HLA-personalized genome | rs71542466 | 32634505 | -39 | 1.00 |
| resting CD4+ memory T cells | present | HLA-personalized genome | rs9276831 | 32832033 | -197567 | 0.01 |
| umbilical cord T cells | Gencord 2, Gutierrez-Arcelus et al, 2013 | reference genome | rs115867015 | 32603487 | 30979 | 0.27 |
| naïve T cells | Blueprint, Chen et al. 2016 | reference genome | rs3129719 | 32661779 | -27313 | 0.05 |
| LCLs | Geuvadis, Aguiar et al., 2018 | HLA-personalized genome | rs9274688 | 32636900 | -2434 | 0.06 |
| LCLs (secondary eQTL) | Geuvadis, Aguiar et al., 2018 | HLA-personalized genome | rs3134978 | 32651477 | -17011 | 0.04 |
| LCLs | Geuvadis, Lappalainen et al, 2013 | reference genome | rs9274660 | 32636434 | -1968 | 0.27 |
| LCLs | Gencord 2, Gutierrez-Arcelus et al, 2013 | reference genome | rs115867015 | 32603487 | 30979 | 0.27 |
| LCLs | GTEx Consortium, 2017 | reference genome | rs17612907 | 32620852 | 13614 | 0.26 |
| macrophages (resting and infected) | Nedelec et al., 2016 | reference genome | rs3135190 | 32667946 | -33480 | 0.06 |
| monocytes | Blueprint, Chen et al. 2016 | reference genome | rs3129758 | 32584625 | 49841 | 0.18 |

